# Disabling a key viral innate immunity antagonist converts measles vaccine into RIG-I–driven cancer immunotherapy

**DOI:** 10.1101/2025.10.03.680227

**Authors:** Aleksandr Barinov, Heidy Vera-Peralta, Joëlle S Nader, Valérie Najburg, Chantal Combredet, Atousa Arbabian, Ségolène Gracias, Phanramphoei N Frantz, Roseline Vibert, Eddy Simard, Marie Coateval, Daniel Pouliquen, Tacien Petithomme, Matthieu Prot, Etienne Simon-Lorière, David Hardy, Sarra Loulizi, Bernadette Brzezicha, Jens Hoffmann, Véronique Riebbels, Jean-François Le Bigot, Marc Grégoire, Jean-François Fonteneau, Anastassia V. Komarova, Nicolas Boisgerault, Frédéric Tangy

## Abstract

Oncolytic viruses can destroy tumors directly or by activating antitumor immunity, but balancing safety with potent immune stimulation remains challenging. Here, we show that deletion of the measles virus C protein, a key viral antagonist of innate immunity, reprograms the live-attenuated vaccine strain into a RIG-I–driven cancer immunotherapy. The resulting virus, MVdeltaC, accumulates defective viral genomes that activate RIG-I/MAVS signaling and trigger robust type I interferon and pro-inflammatory cytokine responses. MVdeltaC kills tumor cells more rapidly and efficiently than the parental virus and induces hallmarks of immunogenic cell death, including HMGB1 release and dendritic cell maturation. Intratumoral administration in immunocompetent mice bearing syngeneic neuroblastoma induced complete tumor regression in 90% of animals and established long-term antitumor memory. Antitumor responses were dependent on CD8⁺ T and NK cells and were further enhanced by anti-CTLA-4 therapy or CD4⁺ T-cell depletion. Prior measles immunization accelerated tumor clearance, indicating vaccine-boosted responses. MVdeltaC also controlled the growth of human mesothelioma, melanoma, and triple-negative breast cancer xenografts and patient-derived tumors in immunodeficient models. These findings establish MVdeltaC as a clinically ready, broad-spectrum immunotherapeutic that links RIG-I activation through defective viral genome generation to elicit potent and durable antitumor immunity.

**IMPACT Statement:** A modified measles virus lacking a viral innate immunity antagonist triggers potent antitumor responses via RIG-I sensing of defective viral genomes, revealing a new strategy for cancer immunotherapy.

## INTRODUCTION

Immune checkpoint inhibition revolutionized cancer treatment with antibodies blocking PD-1, PD-L1, or CTLA-4 to restore T cell activity against tumors. These therapies have shown significant success in treating several malignancies, including melanoma, lung adenocarcinoma, and renal cell carcinoma^1^. However, challenges such as the resistance of non-infiltrated “cold” tumors^2^ and immune-related side effects^3^ limit the clinical efficacy of these treatments. Strategies to turn "cold" tumors into immune-infiltrated "hot" tumors involve enhancing tumor antigen presentation, increasing immune cell recruitment, activating T cells, and boosting type I interferon (IFN-I) signaling to create an inflammatory microenvironment. Immuno-oncolytic viruses (OVs) represent a promising approach, with the prime example of the herpesvirus talimogene laherparepvec (T-VEC) approved for melanoma treatment^4^. More recently, Daiichi Sankyo herpesvirus G47Δ/teserpaturev (Delytact®) received conditional and time-limited marketing approval in Japan as a regenerative medical treatment for malignant glioma^5^, and the non-replicative Ferring’s adenovirus-based gene therapy nadofaragene firadenovec-vncg (Adstiladrin®) was approved for bladder cancer treatment^6^. Other OVs, such as the CG Oncology’s adenovirus Cretostimogene grenadenorepvec or the Replimmune’s herpesviruses Sturlimgene erparepvec and Imlygic talimogene laherparepvec, are being evaluated in phase III clinical trials^7^. These trials focus on both direct tumor cytotoxicity and immune response stimulation and may open the way to OVs combination with checkpoint inhibitors.

Viruses can help establishing adaptive immune memory against the tumor by activating various innate immunity sensors. Among them, retinoic acid-inducible gene I (RIG-I) and melanoma differentiation-associated gene 5 (MDA5) are key cytoplasmic receptors (RIG-I-like receptors, RLRs) that detect viral RNA in cells^8^. RIG-I primarily detects RNA viruses like measles, influenza A virus, and hepatitis C virus^9–11^. These RLRs initiate antiviral responses by triggering IFN-I and cytokine production, promoting immune cell recruitment, including natural killer (NK) and T cells. Beyond their antiviral role, RIG-I and MDA5 serve as tumor suppressors by recognizing abnormal RNA structures from cancer cells and triggering apoptotic pathways and tumor cell death^12^. Their CARD domains interact with mitochondrial antiviral signaling proteins (MAVS), initiating IFN-I release and activating the JAK/STAT pathway, which induces the expression of IFN-stimulated genes (ISGs). Activation of RLR pathways enhances anti-cancer immune responses, and RIG-I agonists are being explored as new therapeutic agents in cancer^8^. MDA5 activation has also been shown to improve checkpoint inhibitor efficacy^13,14^.

Measles virus (MV) is an enveloped, negative sense single-stranded RNA paramyxovirus responsible for measles (Fig. 1a). The attenuated Schwarz strain of MV is commonly used for child vaccination. MV attenuated vaccine strains are also spontaneously oncolytic against numerous types of cancers and have been evaluated in clinical trials for patients with ovarian, glioblastoma, multiple myeloma, mesothelioma, head and neck, breast, and malignant peripheral nerve sheath tumors^15^. MV natural tropism for tumor cells is associated with the overexpression of its entry receptor CD46 on tumor cells, while efficient replication requires defects in IFN-I response^16,17^. We previously observed that MV Schwarz displays spontaneous oncolytic activity against numerous types of cancers, including melanoma^18^, lung and colon adenocarcinomas^19^ and malignant pleural mesothelioma^16,17^. We and others also showed that MV infection induces immunogenic death, thus leading to the maturation of both myeloid and plasmacytoid dendritic cells and promoting tumor antigen cross-presentation^20–23^.

**Figure 1.**
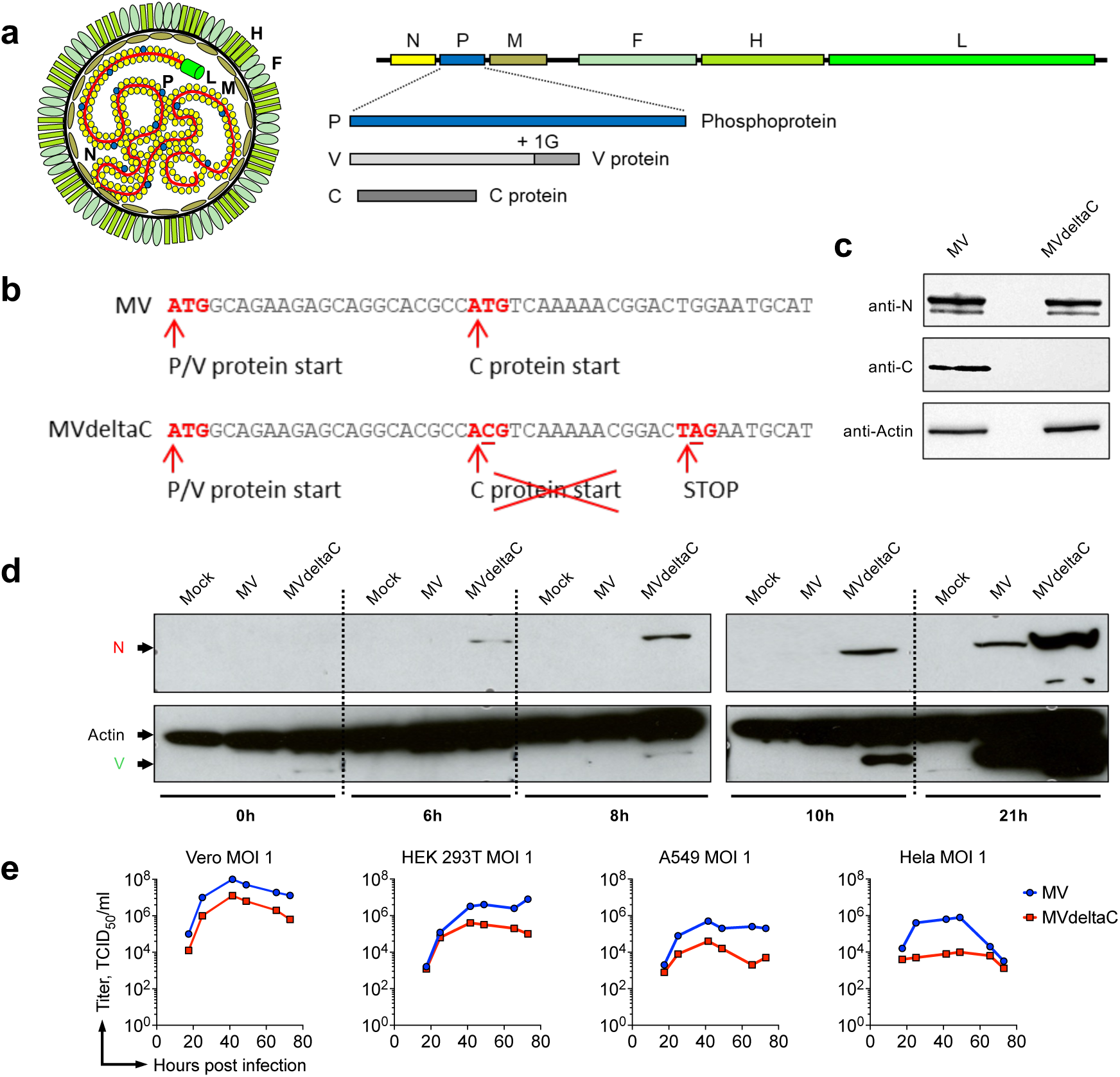
Characterization of MVdeltaC virus. (a) Schematic of the measles viral particle, genome and proteins: nucleoprotein (N), phosphoprotein (P), matrix (M), fusion (F), hemagglutinin (H) and polymerase (L). ORFs for P, V, and C are enlarged. (b) Nucleotide sequence of the P and C ORFs with the mutations (underlined) introduced in MVdeltaC to disrupt C protein expression. (c) Expression of N and C proteins in lysates of Vero cells infected by MV Schwarz or MVdeltaC, analysed by western blot. (d) Kinetics of N and V proteins expression in Vero cell lysates after infection with MV Schwarz or MVdeltaC at MOI 1. (e) Comparative growth kinetics of MV Schwarz (blue) and MVdeltaC (red) in different cell lines infected at MOI 1 and cultured at 37°C. Viruses were harvested from cell lysates at the indicated time points.

To enhance the spontaneous oncolytic activity of attenuated MV and improve its immunostimulatory capacity, we developed MVdeltaC, a C protein–deficient derivative of MV. Measles C protein is a small, cytoplasmic, nonstructural protein produced from an alternative open reading frame within the phosphoprotein (P) gene (Fig. 1a-b). Although lacking enzymatic activity, this virulence factor is incorporated into viral ribonucleocapsids and plays important roles in MV life cycle^24^. It antagonizes host innate immune responses by suppressing IFN-I production^25–27^. It also modulates viral RNA synthesis by acting as a polymerase processivity factor^28,29^. This latter function constrains the formation of defective viral genomes (DVGs), particularly double-stranded RNA (dsRNA). We and others have previously demonstrated that MVdeltaC produces markedly higher levels of DVGs compared to parental MV^9,30^. DVGs, especially 5’ copy-back defective-interfering (DI)-RNAs, are potent RIG-I agonists: their double-stranded structure and 5′ triphosphate ends trigger IFN-I signaling, thereby amplifying antiviral and immune responses^9,31^. In addition, the C protein promotes autophagy, delays apoptosis and supports high-yield viral replication, while mitigating innate immune sensing via mitophagy^32^.

Taken together, these diverse roles make the C protein a strategic target to improve MV immunogenic and oncolytic properties. We hypothesized that C protein deficiency would enhance tumor cell killing by accelerating apoptosis, bypassing autophagy, and increasing immunogenic visibility of the infection. To test this, we investigated the immuno-oncolytic activity and mechanism of action of MVdeltaC both *in vitro* and *in vivo*. Our preclinical findings demonstrate that MVdeltaC is a potent immuno-oncolytic virus against aggressive cancers. With boosted RIG-I activation and IFN-I signaling, MVdeltaC represents a promising strategy for improving the efficacy of immune checkpoint inhibitors and advancing cancer immunotherapy.

## RESULTS

### Generation and characterization of MVdeltaC

MVdeltaC was generated by introducing two mutations into the MV Schwarz genome (GenBank accession no. AF266291.1)^33^: one disrupting the C start codon and another introducing a premature stop codon (Fig. 1b), as previously described^9^. As expected, MVdeltaC-infected cells did not express the C protein (Fig. 1c). Protein expression analysis at early time points revealed that MVdeltaC initiated production of both the structural nucleoprotein (N) and the non-structural V protein significantly earlier (8–10 h post-infection) than parental MV (>21 h), indicating accelerated viral RNA accumulation (Fig. 1d). Although replication in Vero cells was slightly attenuated, MVdeltaC still produced high viral titers (Fig. 1e, Fig S1). To evaluate the genetic stability of the introduced mutations, we performed next-generation sequencing (NGS) of full-length viral genomic RNA after 5 and 10 passages on Vero cells (Fig. S2). The knockout mutations remained intact, which confirmed the genetic stability of MVdeltaC.

### MVdeltaC induces faster tumor cell death than parental MV

We compared the cytotoxic activity of MVdeltaC and the parental MV across a panel of human cancer cell lines using the CellTiter-Glo luminescent viability assay, which measures ATP as an indicator of metabolically active cells. In cervical (HeLa), lung (A549), and mesothelioma (Meso 4, Meso 13 and Meso 163) cell lines, MVdeltaC induced significantly faster and more extensive killing than parental MV (Fig. 2a). Extending this analysis to a panel of bladder, ovarian and hepatocellular carcinoma cell lines showed that 80% of them were efficiently killed by MVdeltaC (viability < 50%, Fig. S3a-b).

**Figure 2.**
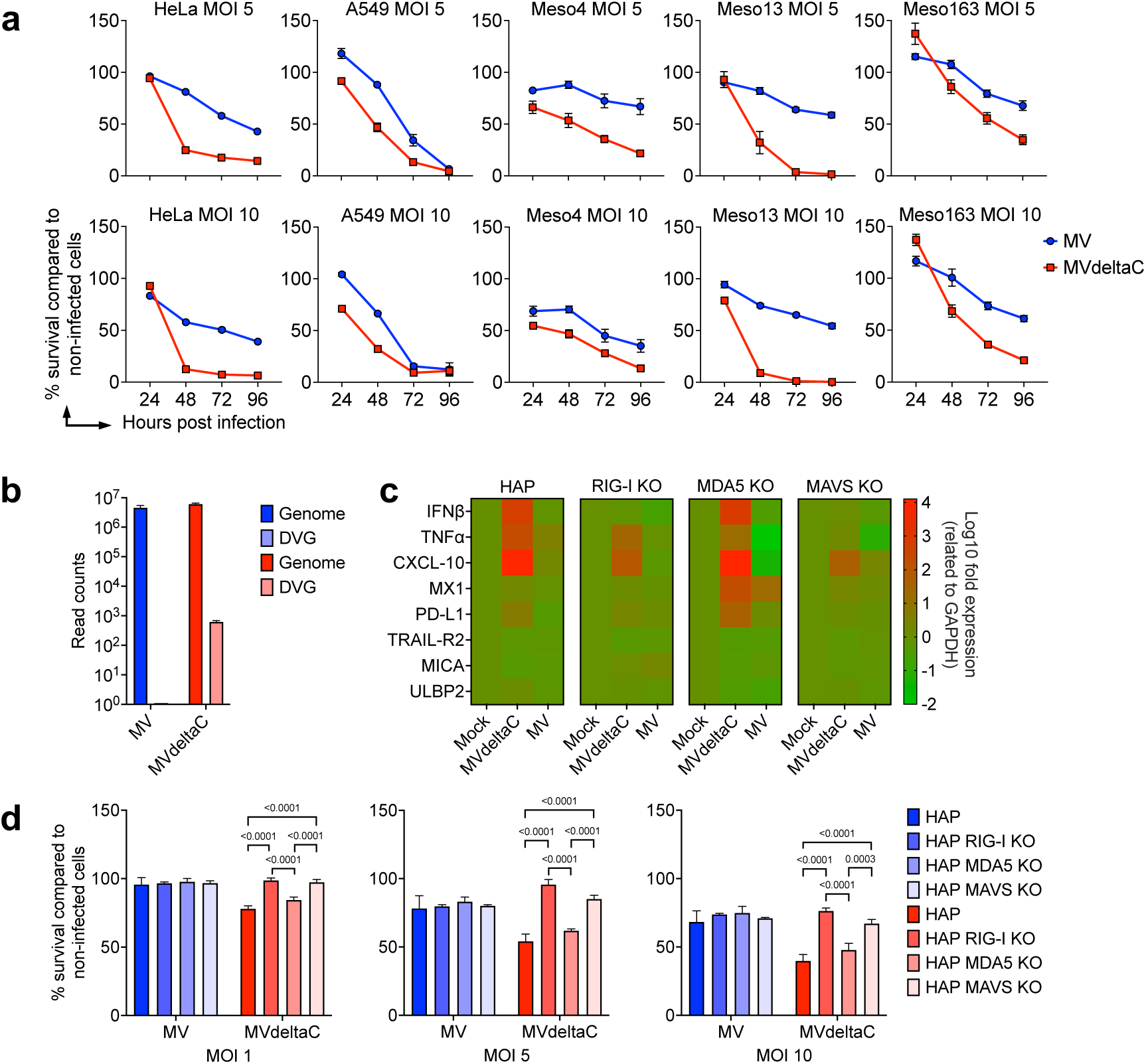
MVdeltaC induces accelerated RIG-I-dependent innate immune activation and killing of cancer cells. (a) Viability of human cancer cell lines infected with MV (blue) or MVdeltaC (red) at increasing MOIs, measured by CellTiter-Glo assay; data normalized to non-infected cells; mean ± SD, n = 3. (b) NGS quantification of DVGs in HAP1 cells infected with MV or MVdeltaC. Bars show total viral reads aligned to MV genome and DVG read counts as detected by DI-tector; mean ± SD, n=3. (c) Innate immunity gene expression profiles in wild-type and knockout HAP1 cells 24 h after infection with MV, MVdeltaC, or PBS (Mock). Heatmap shows log_10_ fold changes of differentially expressed genes; experiment in triplicate, a gradient of colors from green to red represent lowest to highest values, respectively, under all conditions. (d) Viability of wild-type, RIG-I KO, MDA5 KO, and MAVS KO HAP1 cells 24 h after infection with MV or MVdeltaC at MOI 1; means ± SD of three independent experiments; two-way ANOVA test with Šídák’s multiple comparisons test.

### Improved MVdeltaC killing and immunogenicity depends on RIG-I signaling by DVGs

OVs exert their anti-tumor effects not only by direct killing of malignant cells, but also by activating immune pathways in infected cancer cells. We and others have previously shown that, while replicating, MVdeltaC generates high amounts of 5’ copy-back DI dsRNA genomes^9,11,28,30^ that act as RIG-I agonists and strong activators of innate immunity^31,34,35^.

To directly quantify DVG production and dissect their role in enhanced cell killing, we used the human HAP1 cells. These near-haploid, myelogenous leukemia-derived cells possess a functional IFN-I pathway and are available as isogenic knockouts for key innate immune receptors including RIG-I, MDA5, and MAVS, making them suited to study RNA sensing pathways. We first infected wild-type HAP1 cells with either MV or MVdeltaC and performed NGS of total RNA collected from cell lysates 24 h post-infection. Analysis of DVGs was performed using DI-tector, a validated bioinformatic tool for the unbiased detection of DVGs in NGS data^36^. We confirmed that DVGs were produced in MVdeltaC- and not in MV Schwarz-infected cells, consistent with the loss of C protein-mediated suppression of aberrant RNA synthesis (Fig. 2b).

We next examined whether this enhanced DVG production translated into stronger innate immune activation. Gene expression analysis in infected HAP1 cells showed that MVdeltaC induced a markedly stronger innate response than parental MV, with ∼250-fold higher *IFNB1*, 100-fold higher *TNF*, and ∼10,000-fold higher *CXCL10* expression, while other genes such as *MX1*, *CD274* (*PDL1*), *TNFRSF10B* (*TRAILR2*), *MICA*, and *ULBP2* were similarly regulated by both viruses (Fig. 2c). In RIG-I KO cells, MVdeltaC failed to induce *IFNB1*, while *TNF* and *CXCL10* were only modestly upregulated relative to MV. MDA5 KO cells displayed a wild-type-like profile, whereas MAVS KO cells failed to upregulate any of the tested cytokines, except for a modest 10-fold increase in *CXCL10*. In nonmalignant MRC5 fibroblasts, MVdeltaC also triggered significantly stronger *IFNB1*, *TNF*, *CXCL10*, and *MX1* expression compared to parental MV (Fig. S4).

Finally, we tested the functional consequence of this RIG-I signaling on cell killing. HAP1 cells were infected with MVdeltaC or parental MV at different multiplicity of infection (MOI), and cell viability was assessed 24 hours post-infection. While parental MV killed wild-type and KO HAP1 cells at comparable rates, MVdeltaC consistently killed wild-type and MDA5 KO cells more efficiently (Fig. 2d). Importantly, this enhanced killing was abolished in RIG-I KO and MAVS KO cells. Thus, the superior cytotoxic activity of MVdeltaC relies on functional RIG-I and MAVS signaling, while MDA5 does not contribute to this effect.

Together, these results demonstrate that MVdeltaC infection of cancer cells produces abundant RIG-I–stimulating DVGs, which activate MAVS-dependent signaling to drive robust innate immune activation and enhanced cancer cell killing.

### MVdeltaC induces tumor rejection and tumor memory in immunocompetent mice

Because MV is naturally restricted to humans and non-human primates, evaluation of MVdeltaC in immunocompetent mice first required the identification of murine tumor cells permissive to infection. Previous studies showed that the neuroblastoma cell line NS20Y is permissive to MV and, once persistently infected, fails to form tumors in immunocompetent A/J mice^37^. To establish this model, we first confirmed the susceptibility of NS20Y cells to infection using an MV expressing eGFP (MV-GFP). Flow cytometry analysis showed that NS20Y cells were highly permissive to MV compared to MB49 mouse bladder carcinoma cells, used as a non-permissive control (Fig. S5a). Fluorescence microscopy confirmed extensive infection of NS20Y cells, similar to human cancer cells, whereas MB49 cells showed very low infection (Fig. S5b). Infected NS20Y cells also formed large syncytia, indicating a strong fusogenic activity. These results validated the use of the syngeneic NS20Y model in immunocompetent A/J mice to evaluate therapeutic efficacy of MVdeltaC.

Following a preliminary NS20Y cells dose-range study (not shown), mice were subcutaneously (s.c.) grafted with 5 × 10^6^ cells that led to rapid tumor progression. By day 6, all mice developed soft and vascularized tumors ranging from 100 to 250 mm^3^ in volume. Three intratumoral (i.t.) injections of MVdeltaC (2 × 10^6^ TCID_50_) were then performed on days 6, 9, and 12 (Fig. 3a). By day 13, MVdeltaC-treated tumors showed significant volume reduction compared to PBS controls, and immunofluorescence staining of tumor sections with anti-MV N antibody confirmed intratumoral replication of MVdeltaC (Fig. 3b). Treated tumors started to regress between the second and the third MVdeltaC administration and completely disappeared by day 20, leading to survival in 73% of animals (Fig. 3c).

**Figure 3.**
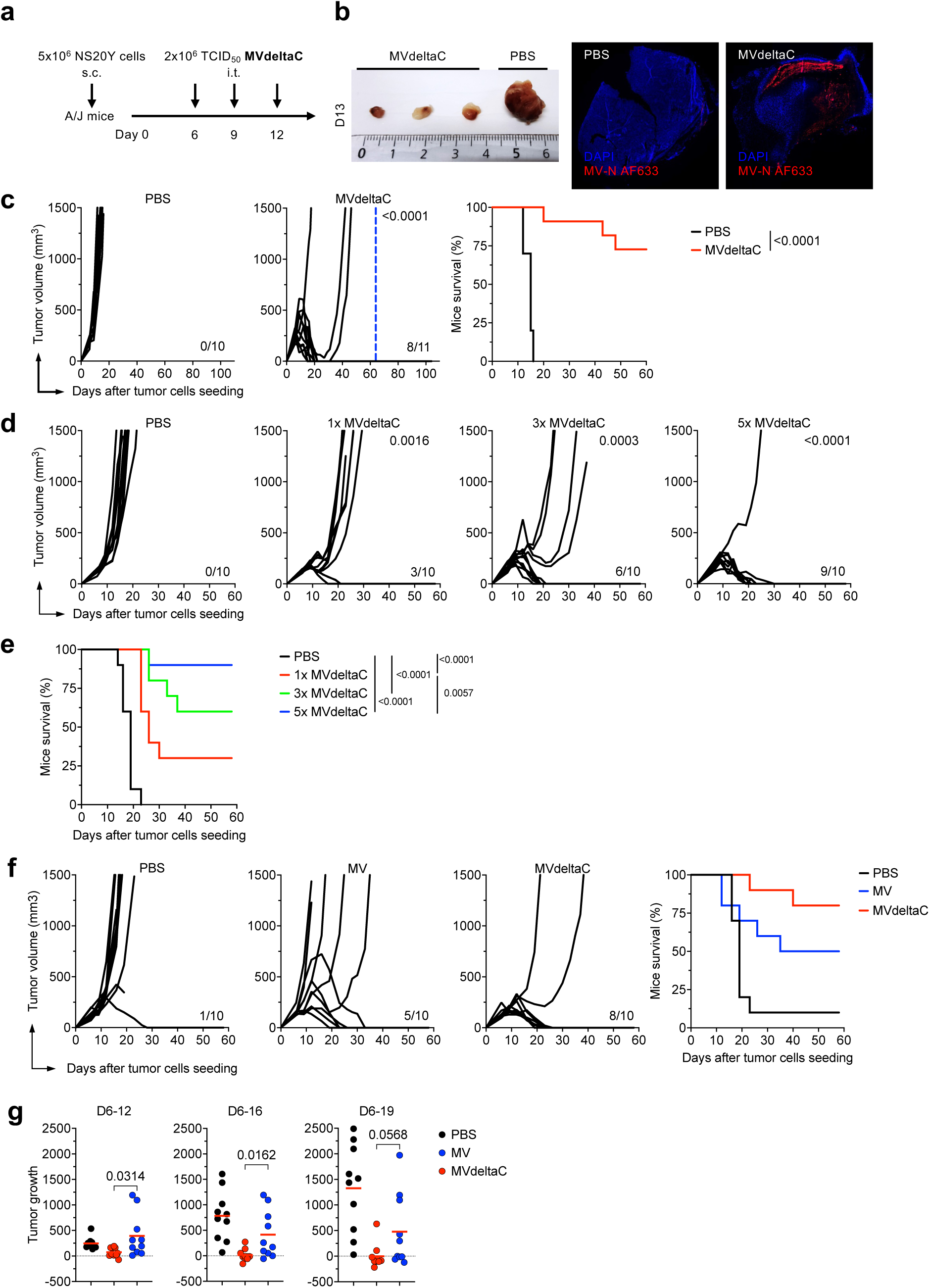
MVdeltaC induces potent NS20Y neuroblastoma tumor rejection in immunocompetent A/J mice. (a) Schematic of the syngeneic NS20Y/AJ model and MVdeltaC treatment schedule. (b) Left, representative tumors excised from the mice on day 13; right, immunofluorescence of tumor cryosections stained for measles nucleoprotein (MV-N, red) and DNA (DAPI, blue). (c) Tumor growth kinetics (black lines, individual mice) and survival after three i.t. injections of MVdeltaC (2 × 10^6^ TCID_50_) or PBS. Blue dotted line indicates the day of tumor rechallenge; n = 10-11 mice per group; MVdeltaC group was compared to PBS-treated group; unpaired one-tailed t test comparing the mean tumor sizes of each group; log-rank test for survival. (d) Tumor growth kinetics after one (1x MVdeltaC, day 6), three (days 6, 9 and 12) or five (days 6, 9, 12, 15 and 19) i.t. injections of MVdeltaC; n = 10 mice per group; each group was compared to PBS-treated group; unpaired one-tailed t test. (e) Kaplan–Meier survival curves for the same groups; log-rank test. (f) Tumor growth kinetics (left) and survival (right) of mice treated with three i.t. injections of parental MV, MVdeltaC (at the same dose of 0.5 × 10⁶ TCID₅₀), or PBS over a 2-month observation period; n = 10 mice per group; log-rank test for survival. (g) Comparison of tumor control by MVdeltaC (red) and parental MV (blue). Tumor growth inhibition calculated as the change in tumor volume between treatment initiation (day 6) and study days 12, 16, or 19; n = 10 mice per group; unpaired two-tailed t-test.

To determine whether MVdeltaC treatment elicited tumor-specific immune memory, cured mice were re-challenged at day 65 with the same number of NS20Y cells injected into the contralateral flank (blue line in Fig. 3c). None of the mice developed tumors within 40 days post re-challenge, demonstrating the induction of durable, tumor-specific adaptive immune memory.

We next tested the impact of administration frequencies. Mice received one, three, or five i.t. injections of MVdeltaC (Fig. 3d-e). Five i.t. injections resulted in complete tumor regression in 90% of mice, indicating that increasing the frequency of injections significantly enhances the efficacy.

### MVdeltaC demonstrates superior antitumor efficacy over the parental MV in immunocompetent mice

To directly compare MVdeltaC with the parental MV in the same mouse model, we assessed tumor control by both viruses over a 2-month period (Fig. 3f). Three i.t. injections of MVdeltaC induced complete tumor regression in 80% of animals, whereas the parental MV achieved complete response in 50%. To quantitatively evaluate treatment efficacy, tumor growth inhibition (TGI) was calculated for each animal as the difference between tumor volume on a given day and tumor volume at treatment initiation (day 6). Positive TGI values reflect tumor progression relative to baseline, while values near zero or negative indicate stabilization or regression. All animals, including those sacrificed early due to tumor burden, were included in the analysis using their final recorded tumor volumes. TGI analysis revealed significantly greater tumor inhibition in the MVdeltaC group compared with the parental MV on days 12 and 16 (p = 0.0314 and p = 0.0162, respectively; Fig. 3g and Table 1). At day 12, MVdeltaC treatment reduced mean tumor growth by approximately 70% (75 mm^3^ vs. 246 mm^3^ in PBS controls), while parental MV showed no significant effect (394 mm^3^). By day 16, tumor growth was almost completely inhibited in 80% of animals (mean 14 mm^3^ vs. 783 mm^3^ in PBS; 98% inhibition), whereas parental MV achieved only 47% inhibition (415 mm^3^). By day 19, MVdeltaC maintained durable tumor control (mean TGI = –3 mm^3^), indicating sustained regression, while MV-treated animals continued to show tumor progression (mean TGI = 478 mm^3^; p = 0.0568).

**Table 1.**
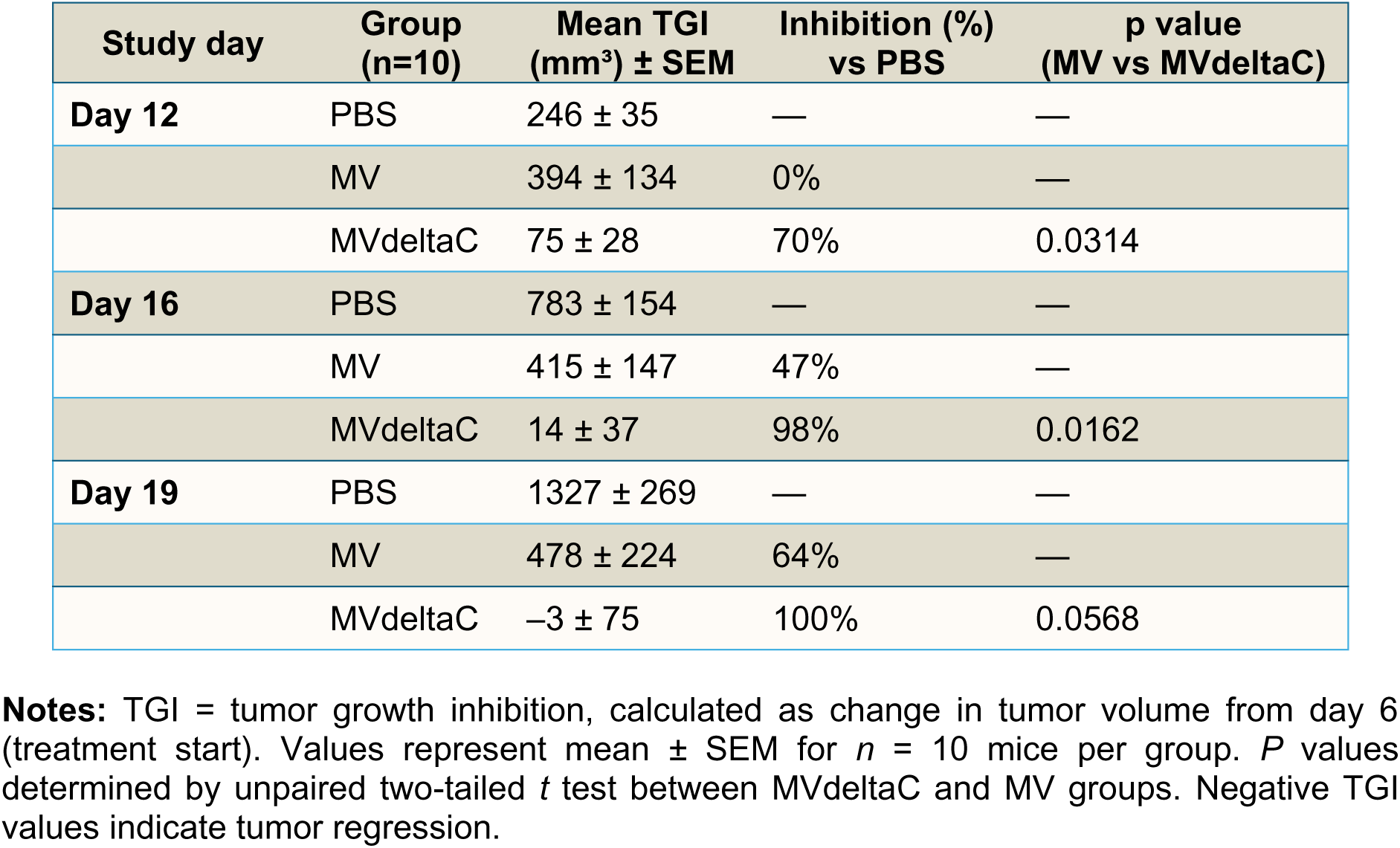
| Tumor growth inhibition analysis of NS20Y tumors following i.t. MV or MVdeltaC treatment

Collectively, these findings demonstrate that MVdeltaC exhibits potent and durable antitumor activity in immunocompetent mice, markedly surpassing that of the parental MV. Deletion of the C protein provides a clear therapeutic advantage *in vivo*. Intratumoral administration triggered virus replication and complete tumor eradication in most treated animals, and the cured mice developed durable, tumor-specific immune memory.

### Lymphocytes play a major role in the therapeutic activity of MVdeltaC

To assess the role of adaptive immune cells in MVdeltaC efficacy, we first examined tumor-infiltrating lymphocytes (TILs) by immunohistochemistry of regressing NS20Y tumors collected at day 13. Abundant infiltration of T cells was observed in MVdeltaC-treated tumors (Fig. 4a). Both CD4⁺ and CD8⁺ T cells accumulated around the tumors and were homogeneously distributed throughout the tumor tissue, thus demonstrating that MVdeltaC treatment converted the tumor microenvironment into an immunologically active “hot” state.

**Figure 4.**
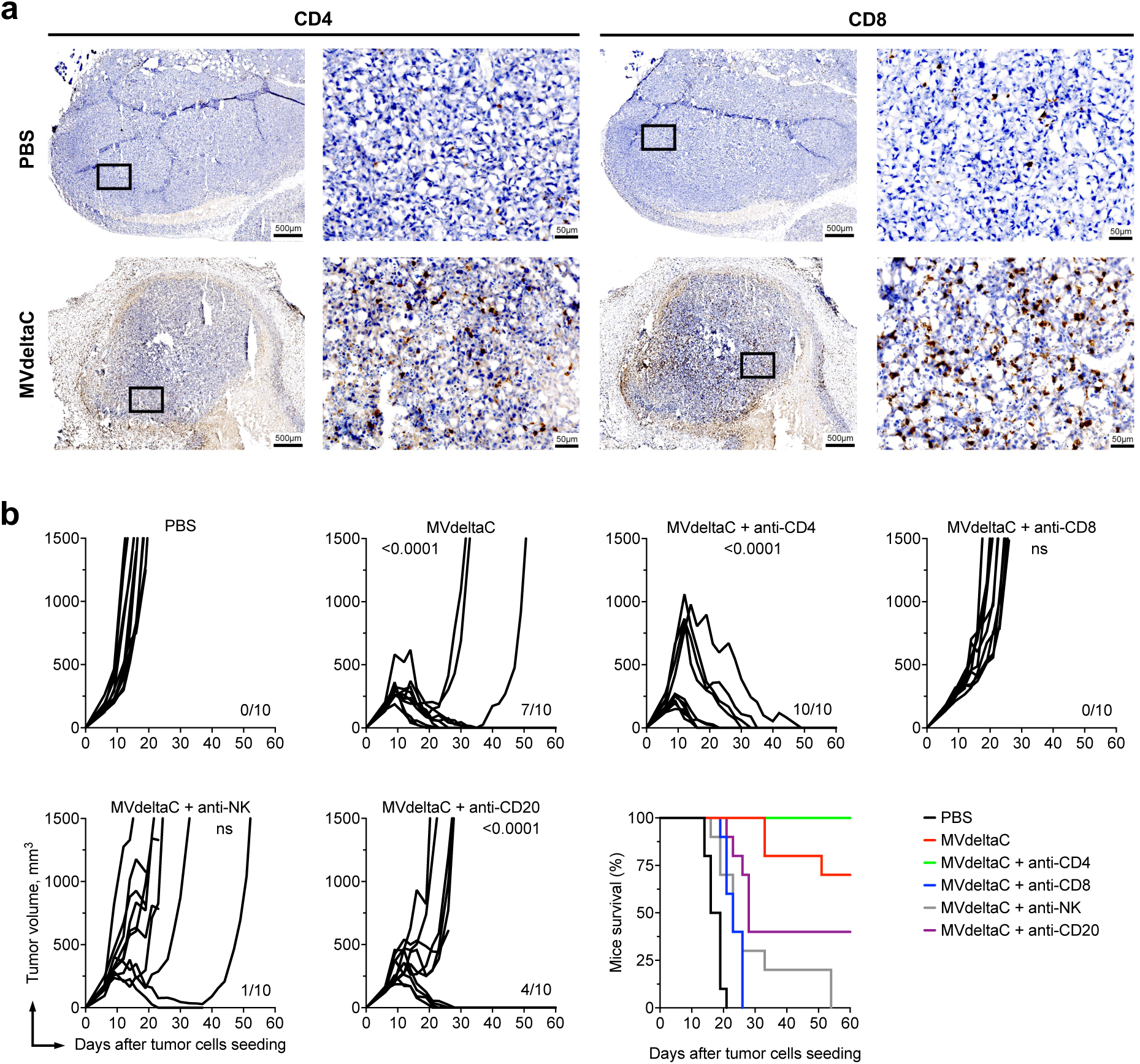
MVdeltaC induces massive T cells infiltration into tumors, and rejection efficacy relies on a functional immune system. (a) Immunohistochemistry of NS20Y tumors 24 h after the third i.t. injection of MVdeltaC, stained for CD4^+^ and CD8^+^ T cells. (b) Tumor growth and survival in mice depleted of CD4^+^, CD8^+^, B cells or NK cells during MVdeltaC therapy. Depletion was maintained by weekly antibody administration until day 49. Tumor therapy with MVdeltaC was performed on days 6, 9 and 12; *n =* 10 mice per group; each group was compared to PBS-treated group; unpaired one-tailed t test; ns, not significant.

We next investigated which immune cell subsets were involved in MVdeltaC immuno-oncolytic activity by selectively depleting CD4^+^, CD8^+^ T cells, B cells, or NK cells in A/J mice grafted with NS20Y neuroblastoma tumors. Flow cytometry confirmed efficient depletion of the targeted immune cell subsets (Fig. S6). Depletion of CD4^+^ T cells enhanced the therapeutic efficacy of MVdeltaC (Fig 4b). All the mice in this group achieved complete tumor rejection, compared to a 70% complete response rate in non-depleted controls. On day 12, at the third MVdeltaC injection, tumor volumes in the CD4^+^ T cell-depleted group were heterogeneous: four mice exhibited very large tumors (mean volume = 887mm^3^), whereas six mice had smaller tumors (mean volume = 152mm^3^, Fig. 4b). Mice with smaller tumors cleared them more rapidly (by days 16-23), while mice with larger tumors achieved complete regression later (days 30–49). These results demonstrate that even very large tumors can be eliminated by MVdeltaC in the absence of CD4^+^ T cells. All mice remained tumor-free up to 100 days. In contrast, depletion of CD8^+^ T cells and NK cells abrogated the therapeutic efficacy of MVdeltaC (Fig. 4b). Tumor growth kinetics in the absence of CD8^+^ T cells was similar to that observed in PBS-treated controls. NK cell depletion also largely eliminated therapeutic benefit. B cell depletion also impaired the efficacy of MVdeltaC treatment, albeit to a lesser extent than CD8^+^ T cell or NK cell depletion. Only 40% of B cell-depleted mice exhibited complete tumor response, indicating that while B cells contribute to the therapeutic outcome, they are not as crucial as CD8^+^ or NK cells.

Taken together, these findings indicate that CD4^+^ and CD8^+^ T cells play critical but opposite roles in MVdeltaC therapy. Depletion of CD4^+^ T cells enhances the therapeutic efficacy of MVdeltaC, likely *via* Treg removal, whereas the absence of cytotoxic CD8^+^ T cells and NK cells completely abolishes the immuno-oncolytic efficacy of MVdeltaC, resulting in progressive disease. Intratumoral administration of MVdeltaC thus promotes significant infiltration of both CD4⁺ and CD8⁺ T cells into the tumor microenvironment, transforming it into an immunologically “hot” state and supporting tumor rejection through the induction of adaptive antitumor immunity.

### Preexisting measles immunity does not impair MVdeltaC efficacy

Since most humans are immunized against measles through vaccination or infection, we assessed whether this preexisting immunity affects MVdeltaC therapy. We immunized mice with the parental MV (MV-immunized group) or PBS (MV-naïve controls). Thirty-eight days later, high MV-specific IgG antibody titers confirmed successful immunization (Fig. S7). Mice were then grafted s.c. with NS20Y tumors and treated with MVdeltaC starting on day 6. Tumor responses were comparable: complete response rates were 77.7% in MV-naïve and 71.4% in MV-immunized mice (Fig. 5a), with no significant survival differences (Fig. 5b). Notably, tumor clearance was faster in immunized mice (median 21 vs. 33 days), though not statistically significant (Fig. 5c). These results suggest that preexisting measles immunity does not diminish, and may even accelerate, MVdeltaC-mediated tumor clearance.

**Figure 5.**
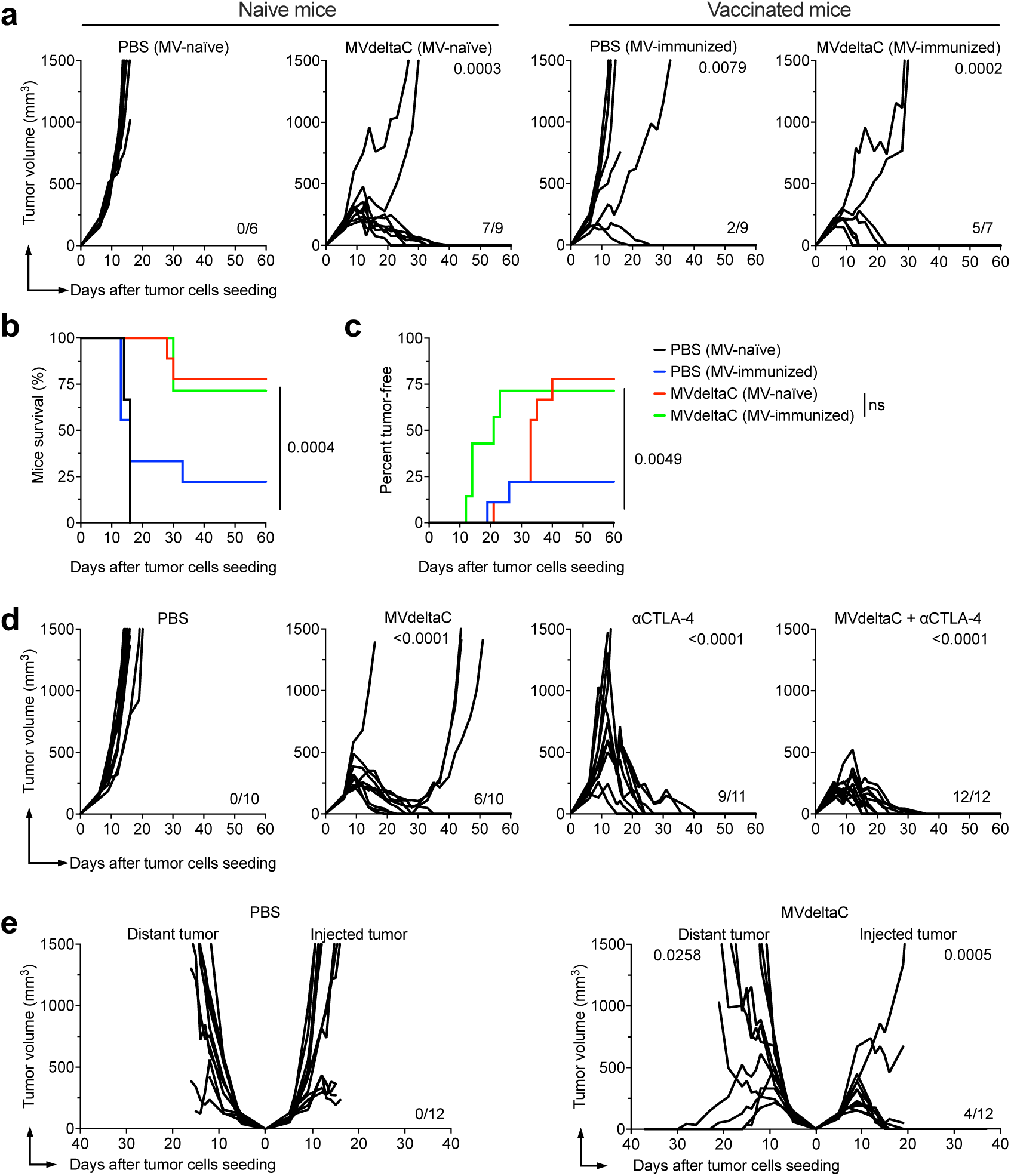
MVdeltaC therapy is not affected by preexisting measles immunity, is enhanced by anti-CTLA-4 therapy and exhibits abscopal effect. (a) NS20Y tumor growth in MV-naïve or measles pre-vaccinated mice treated i.t. with MVdeltaC or PBS on days 6, 9, 12 and 19 after tumor cells seeding. (b) Survival rates, and (c) percentage of tumor-free animals in naïve or pre-vaccinated mice; *n =* 6-9 mice per group; each group was compared to MV-naïve PBS-treated group, unpaired one-tailed t test; log-rank test for survival. (d) Tumor growth in mice treated with three i.t. injections of MVdeltaC and/or four i.p. injections of anti-CTLA-4 (αCTLA-4) antibody on days 6, 9, 12 and 15; n = 10-12 mice per group; each group was compared to PBS group (unpaired one-tailed t test). (e) Growth of injected and contralateral uninjected (distant) tumors in mice treated with MVdeltaC in single tumor; n = 12 mice per group; MVdeltaC-injected and distant tumors were compared to their corresponding PBS controls; unpaired one-tailed t test.

### Synergistic activity of MVdeltaC with anti-CTLA4 immune checkpoint inhibitor and abscopal effect

Considering the immunotherapeutic potential of MVdeltaC, we next tested MVdeltaC in combination with immune checkpoint blockade targeting CTLA-4 (Fig. 5d). In NS20Y-bearing mice, MVdeltaC monotherapy resulted in a complete response in 60% of treated mice, anti-CTLA-4 monotherapy led to complete tumor clearance in 82% of animals, whereas combination therapy of MVdeltaC and anti-CTLA-4 resulted in complete tumor regression in 100% of mice, demonstrating the additive or synergistic effects of both therapies. This finding highlights the translational potential of MVdeltaC as a promising component for combination immunotherapy regimens.

Finally, we assessed whether MVdeltaC could elicit systemic antitumor responses against untreated contralateral tumors in a double-tumor model. Although the NS20Y model in A/J mice is very aggressive – tumors grow very fast compared to the establishment of the adaptive CD8 responses – a notable antitumor response was observed after treatment with MVdeltaC. Specifically, 10 out of 12 mice treated with MVdeltaC showed a decrease in tumor volume in the injected tumors. Among these, 4 mice (40%) achieved complete regression of both injected and non-injected tumors (Fig. 5e). This abscopal activity indicates that MVdeltaC can generate systemic antitumor immunity capable of controlling distant lesions.

### Translating the immuno-oncolytic activity of MVdeltaC in patient-derived cancer cells

With the aim to translate the development of MVdeltaC to human cancers, we analyzed its oncolytic capacity against a collection of human pleural mesothelioma (PM) cells. We previously studied the varying susceptibility of human PM cells to MV replication^16^. Here, using fluorescent microscopy, we analyzed the infection of twelve PM cell lines by MVdeltaC-eGFP (Fig. 6a-b and S8 + S9). As expected, healthy mesothelial cells (MES-F and LP-9) (Fig. 6a-b), as well as fibroblasts and endothelial cells (Fig. S8a) were resistant to both MV and MVdeltaC, confirming the tumor selectivity of oncolytic MV. Meso 4 cells known to be resistant to parental MV at low MOI were also resistant to MVdeltaC replication, whereas most PM cell lines were permissive (Fig. 6 a-b, S8b, S9b). GFP expression intensity was lower in MVdeltaC-infected cells compared to parental MV (Fig. 6a + S9), which confirmed lower viral replication, even if the proportions of infected cells were similar between both viruses (Fig. 6b).

**Figure 6.**
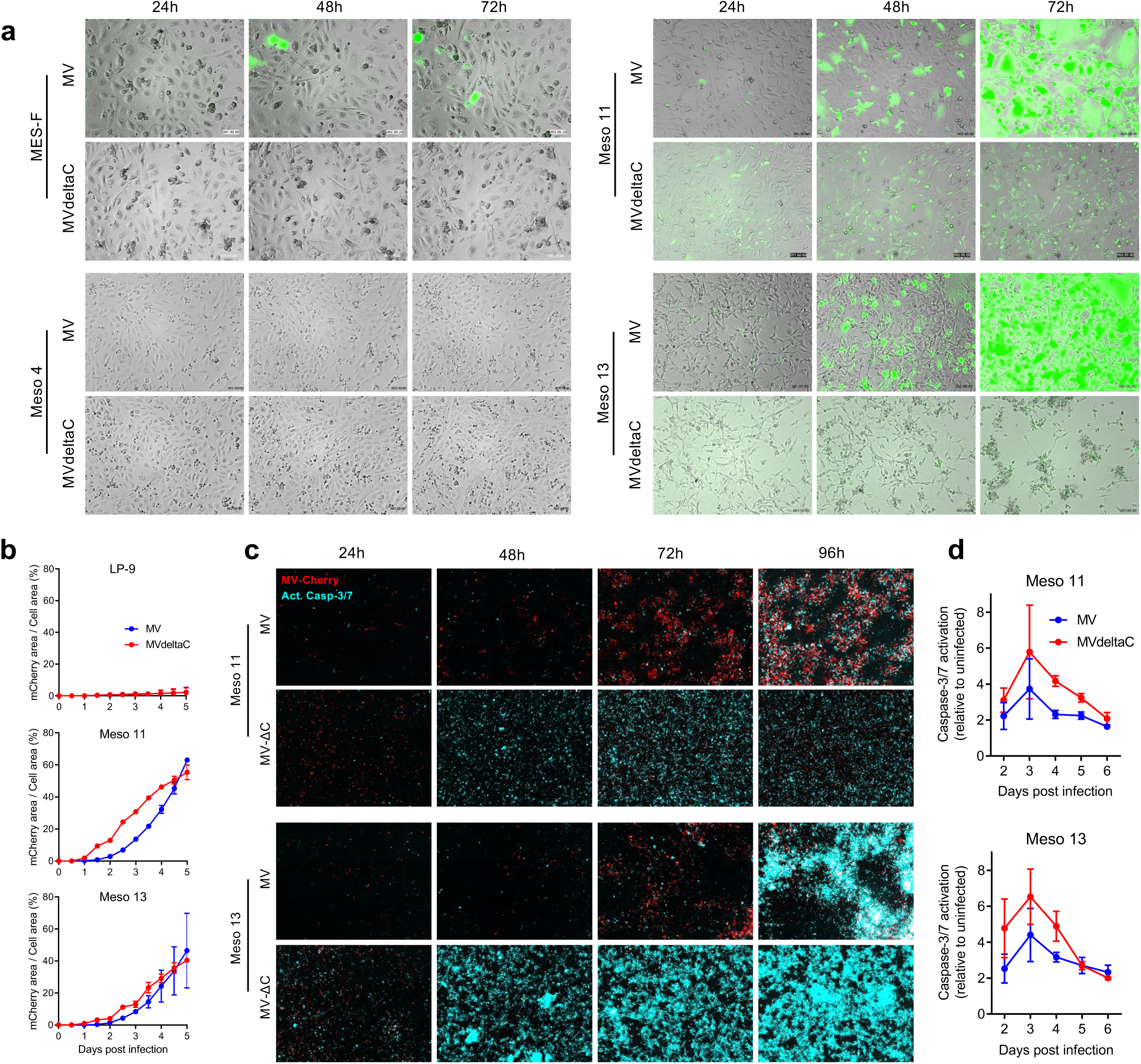
Oncolytic activity of MVdeltaC in human mesothelioma cells. (a) Time-lapse microscopy of healthy mesothelial (MES-F) and mesothelioma (Meso 4, Meso 11, Meso 13) cells infected with MV-eGFP or MVdeltaC-eGFP at MOI 1. (b) Quantification of virus propagation in normal mesothelial (LP-9) and mesothelioma (Meso 11, Meso 13) cells infected with either MV-mCherry or MVdeltaC-mCherry. (c) Caspase-3/7 activation (NucView probe, cyan) detected by fluorescent microscopy in mesothelioma cells infected with either MV-mCherry or MVdeltaC-mCherry (red). (d) Caspase-3/7 activity measured by luminescence (Caspase-3/7-Glo). Data are mean ± SD of at least three independent experiments.

To examine cell death induction, we analyzed caspase-3/7 activation over time by fluorescence microscopy (Fig. 6c, S10d) and a luminescence bioassay (Fig. 6d, S10a-c). MVdeltaC triggered apoptosis earlier than parental MV, with robust caspase-3/7 activation observed as early as 48 h post-infection in sensitive lines, compared to 96 h for MV (Fig. 6c). Quantification of caspase-3/7 activation in twelve human MPM cell lines over 6 days confirmed that MVdeltaC elicited enhanced apoptosis in several models (Fig. 6d, S10a-d). Meso 11, Meso 13, Meso 76 and Meso 225 showed enhanced oncolytic activity of MVdeltaC over MV, whereas Meso 34, Meso 47 and Meso 163 showed similar activation of caspase-3/7 between MV and MVdeltaC (Fig. 6d, S10a-b). Interestingly, two cell lines, Meso 35 and Meso 56, appeared to be resistant to MVdeltaC but not to MV (Fig. S10b). The three cell lines resistant to MV (Meso 4, Meso 150, Meso 173) were also resistant to MVdeltaC (Fig. S10c-d). Notably, while at standard MOI of 1 Meso 4 was resistant to replication and killing, this cell line was nevertheless killed when using higher MOIs of 5 and 10 (Fig. 2a).

Together, these results demonstrate that MVdeltaC efficiently infects and kills a large majority of patient-derived PM cell lines by inducing faster apoptotic death than the parental MV, while retaining selectivity for malignant over normal mesothelial and stromal cells.

### MVdeltaC accumulates viral dsRNA, triggers the release of danger signals and activates dendritic cells

We next determined whether MVdeltaC infection resulted in the accumulation of viral dsRNA in the cytoplasm of infected human PM cells. Indeed, confocal microscopy revealed that dsRNAs were present in MVdeltaC-infected cells but absent in parental MV-infected or uninfected cells (Fig. 7a).

**Figure 7.**
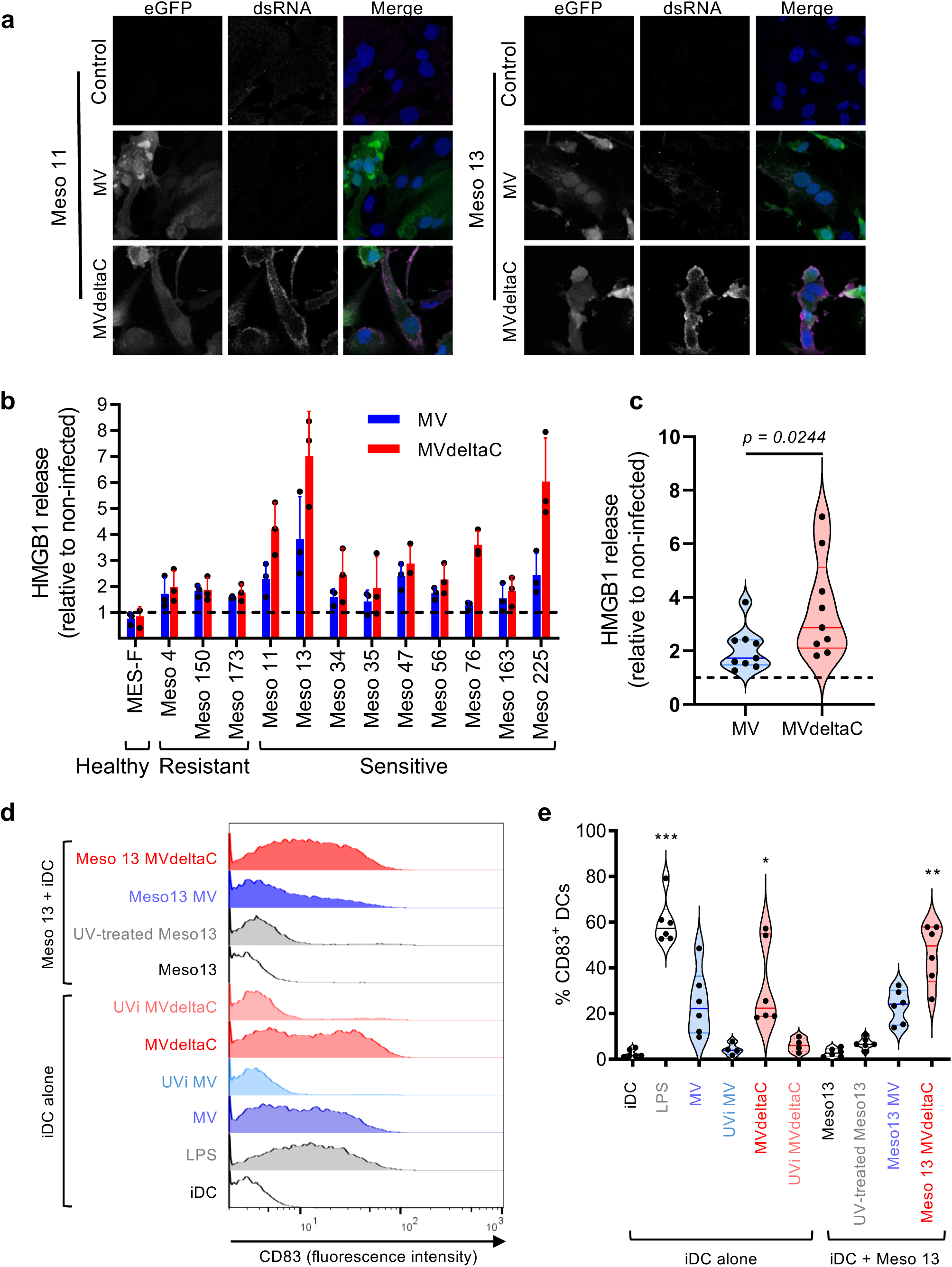
**MVdeltaC induces danger signal production by tumor cells and promotes dendritic cell maturation. (**a) Confocal microscopy of dsRNA (purple) in mesothelioma cells infected for 48h with either MV-eGFP or MVdeltaC-eGFP (MOI = 1, green). DNA is labeled with Hoechst (blue). (b) HMGB1 release in supernatants of healthy mesothelial MES-F cells and mesothelioma cells measured by ELISA 72 h after infection with MV Schwarz or MVdeltaC (MOI=1). Data represent means ± SD of three independent experiments. (c) Comparison of HMGB1 release between the MV and MVdeltaC conditions for MV-sensitive cell lines presented in (b); two-tailed Mann-Whitney test. (d-e) Immature human dendritic cells (iDCs) cultured for 20h in presence of LPS (1µg/mL), MV Schwarz or MVdeltaC (2 × 10^5^) previously UV-inactivated or not, or Meso13 mesothelioma cells previously exposed to UV-B (10J/cm²) or infected for 72h with MV Schwarz or MVdeltaC. HLA-DR^+^ DCs were then analyzed by flow cytometry for CD83 surface expression as shown for one experiment (d) or for 6 independent experiments (e); Kruskal-Wallis test.

Immunogenic cell death (ICD) is characterized by the release of molecules of danger-associated molecular patterns (DAMPs), such as ATP, calreticulin, or HMGB1, which act as signals to activate DCs and prime adaptive immune responses. Both MV and MVdeltaC infection induced HMGB1 release during tumor cell death, with higher amounts being released from MVdeltaC-infected tumor cells (Fig. 7b-c). We then tested the functional immunogenicity of MVdeltaC-induced cell death by evaluating its ability to promote the maturation of human monocyte-derived immature dendritic cells (iDCs) from six healthy donors, either directly or after infection of PM cells (Fig. 7d-e). As expected, direct exposure of iDCs to MV or MVdeltaC induced the expression of the CD83 maturation marker on their surface. This effect was completely abrogated when both viruses were previously UV-inactivated, indicating that viral replication is necessary to induce DC maturation. We previously showed that infection of Meso 13 cells by parental MV Schwarz promoted iDC maturation^20^. Infection of Meso13 cells with MVdeltaC at the same MOI further enhanced this activation, thereby showing superior activity of MVdeltaC at inducing ICD and activating antigen-presenting cells (Fig.7e). Control Meso 13 cells left untreated or induced into apoptosis by ultraviolet radiation did not trigger DC maturation.

Altogether, these results demonstrate that MVdeltaC infection of patient-derived tumor cells leads to dsRNA accumulation, danger signal release, and efficient induction of DC maturation — hallmarks of ICD that support initiation of antitumor immune responses.

### MVdeltaC controls human cancers in orthotopic and PDX models

To assess MVdeltaC oncolytic activity *in vivo* against patient-derived tumor cells, we first performed orthotopic xenograft experiments in NOD/SCID mice. These animals lack functional T and B lymphocytes, have very low NK cell activity and an absence of hemolytic complement, while macrophages and DCs are present but functionally impaired. Of the seven MVdeltaC-sensitive human PM cell lines tested, only Meso 34 and Meso 163 successfully established tumors in the peritoneal cavity. Three weeks post-grafting for Meso 163 and two months for Meso 34, mice received a single low-dose i.p. injection of MVdeltaC (2 × 10^5^ TCID_50_) or control PBS. Two weeks later, tumor burden was significantly reduced in the MVdeltaC-treated group for both cell lines compared to control animals (Fig. 8a). MVdeltaC treatment efficiently reduced the number and the size of tumors in the peritoneum (Fig. 8b). Histology analysis revealed that tumor density was strongly decreased in animals treated with MVdeltaC compared to control mice (Fig. 8c).

**Figure 8.**
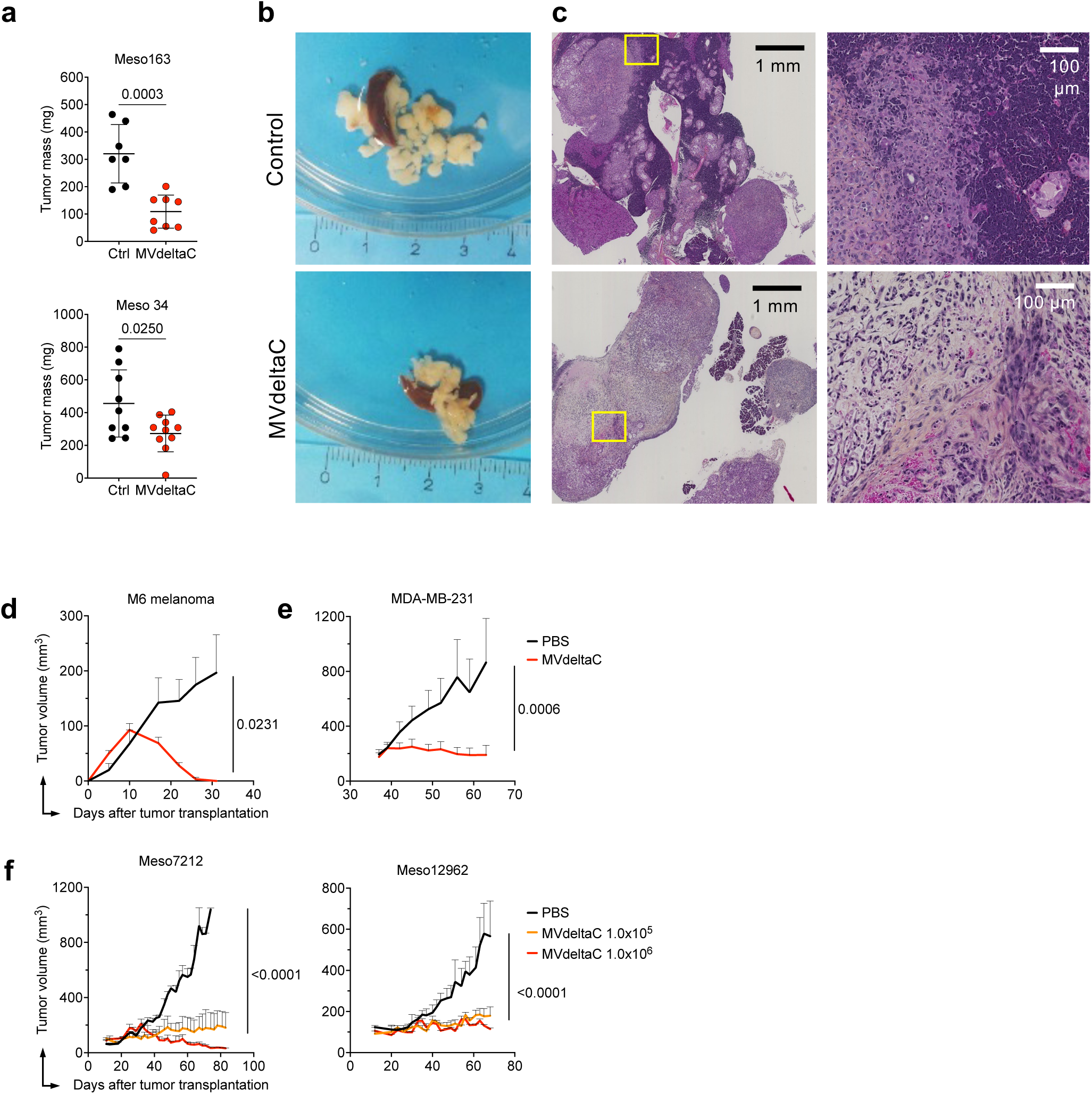
MVdeltaC controls human tumor growth in immunodeficient mouse models. (a) Meso 163 (upper panel) and Meso 34 (lower panel) tumor weights in NOD/SCID mice two weeks after single i.p. injection of MVdeltaC; means ± SD, n=7 - 10 mice per group; unpaired two-tailed t test. (b) Representative images of Meso 34 tumor nodules from treated and control animals. (c) Hematoxylin-phloxine-saffron staining of the peritoneum with detailed view of the tumor area (right). (d) Human M6 melanoma tumor growth in NOD:SCID mice treated with two i.t. injections of MVdeltaC on days 10 and 17; means ± SEM, n = 4 mice per group; one-tailed t test. (e) Triple-negative breast cancer MDA-MB-231 tumor growth in NMRI nu/nu mice after weekly i.t. injections of MVdeltaC. Therapy started on day 38, for a total of eight bi-weekly injections; means ± SEM, n=6 mice per group; unpaired one-tailed t test. (f) mesothelioma PDX tumor growth in NMRI nu/nu mice after weekly MVdeltaC therapy at two doses; means ± SEM, n = 3 mice per group; each group was compared to PBS group (unpaired one-tailed t test).

To evaluate whether the oncolytic effect of MVdeltaC extends beyond PM, we next tested human melanoma and triple-negative breast cancer (TNBC) xenografts. In NOD/SCID mice bearing subcutaneous M6 human melanoma xenografts, i.t. administration of MVdeltaC significantly reduced tumor growth compared to PBS-treated controls (Fig. 8d). Similarly, in NMRI nu/nu mice (deficient in T lymphocytes but retaining NK cells, DCs, macrophages, and an intact complement activity) s.c. xenografts of the human TNBC cell line MDA-MB-231 were efficiently controlled by weekly i.t. injections of MVdeltaC (Fig. 8e). As these mice lack T cell memory, repeated virus administration was required throughout the study.

Tumor models based on established tumor cell lines have the disadvantage of low heterogeneity^38^. Therefore, we next evaluated MVdeltaC in patient-derived mesothelioma xenografts (PDX) serially passaged in NMRI nu/nu mice. Four PDX models established from patients with pleural or peritoneal mesothelioma were selected: Meso7212, Meso12962, Meso13852B and Meso14275^39,40^. When tumors reached ∼100 to 250 mm^3^, mice received weekly i.t. injections of MVdeltaC. Two doses of MVdeltaC were tested: 1 × 10^5^ and 1 × 10^6^ TCID_50_ per injection. MVdeltaC treatment significantly inhibited tumor growth across all four PDX models at both doses (Fig. 8e and Fig.S11). According to the modified RECIST criteria^41^, treatment responses in Meso7212 and Meso12962 tumors were stable disease, whereas treatment with the higher dose induced partial response (Fig 8f). Therapy was well tolerated without side effects nor body weight loss (not shown).

Together, these findings establish MVdeltaC as a broad-spectrum oncolytic virus with potent activity across mesothelioma, melanoma, and TNBC models, highlighting its translational potential for clinical development in multiple solid tumor indications.

## DISCUSSION

In this study, we investigated the immuno-oncolytic potential of MVdeltaC, a modified measles vaccine virus in which the nonstructural regulatory protein C, which plays a central role in antagonizing the host innate immune response, was deleted. We observed that MVdeltaC kills tumor cells more rapidly and efficiently than the parental virus *in vitro* and *in vivo*. This enhanced anticancer activity depends on RIG-I activation by DVGs generated by the modified virus, leading to rapid and strong expression of pro-inflammatory cytokines and chemokines. In consequence, both the virus and infected tumor cells become more visible to the host immune system. Such activity promotes strong DC maturation and the recruitment of immune cell (CD8+ T cells, NK cells) into the tumor that participate to the killing of tumor cells.

MVdeltaC demonstrated potent anti-cancer effects *in vitro*, inducing rapid tumor cell death in over 40 human cancer cell lines, including mesothelioma, lung adenocarcinoma, bladder cancer, ovarian cancer, hepatocarcinoma, and cervical cancer. MVdeltaC killed cells more rapidly and more extensively than parental MV and remained unable to spread in non-malignant cells, as deletion of the C protein further attenuated the virus. Tumor cell death was associated with the release of large quantities of danger signals, including cellular (HMGB1) and viral (dsRNA) components. Extremely high levels of *IFNB1*, *TNF* and *CXCL10* were induced in dying cancer cells, with as much as 10,000 times more *CXCL10* expression than after infection by standard MV. Notably, the strong induction of *CXCL10* suggests a mechanism by which MVdeltaC could promote the recruitment of immune cells into the tumor microenvironment.

*In vivo*, MVdeltaC effectively inhibited tumor growth in immunodeficient mouse models. In NOD/SCID mice grafted orthotopically with human mesothelioma cells, a single, low-dose intraperitoneal injection significantly reduced tumor mass within two weeks. Similar efficacy was confirmed in mice bearing TNBC and human melanoma cells, as well as against mesothelioma PDX. In immunocompetent A/J mice grafted with syngeneic neuroblastoma cells, three i.t. injections of MVdeltaC led to complete tumor rejection in 70% of animals, and five injections in 90% of them. Treatment also generated immune memory, as surviving animals rejected tumor re-challenge. Regressing tumors exhibited massive infiltration of CD8+ and CD4+ T cells, and depletion studies revealed that MVdeltaC efficacy was dependent on CD8+ T lymphocytes and NK cells, while depletion of CD4+ T cells or a combination with anti-CTLA-4 enhanced efficacy. Even though the NS20Y model is very aggressive and we had to treat large tumors, an abscopal effect was also observed likely due to CD8+ T cell and NK cell activity. While CD8^+^ T cells are essential for long-term tumor control, their efficacy is significantly diminished in the absence of NK cells. NK cells likely play a crucial role in the early recognition and killing of MVdeltaC-infected tumor cells, which may be necessary to initiate an effective CD8^+^ T cell response. Without this early innate activation, CD8⁺ T cells may receive insufficient priming and fail to control rapidly growing tumors. Importantly, prior measles immunity did not hinder efficacy but rather accelerated anti-tumor responses. This is a major finding considering that a vast majority of the population is immune to MV and that the actual clinical impact of both cellular and humoral immune responses directed against OVs is still debated.

This work demonstrates that by introducing two mutations into the 15,894 nucleotides of the measles vaccine virus genome, we generated a highly immuno-oncolytic virus with a greater capacity to kill cancer cells and to elicit strong immunostimulatory responses prone to attract and activate immune cells into the tumor microenvironment. Unmodified MV live-attenuated strains are naturally oncolytic against many malignancies^15^ and are currently evaluated clinically for patients with different types of cancers^42^. Modifications of oncolytic MV were first mainly focused on increasing its safety, for example by inserting the *IFNB1* gene into the MV genome or by retargeting the virus toward alternative tumor-specific receptors. Considering the advances in cancer immunotherapy, most of the modification of OVs now focus on increasing their immuno-activating properties. Particularly with MV, a number of modifications have been proposed, such as inserting GM-CSF, IL12, or anti-PD1 and anti-CTLA4 single chain antibodies sequences into the virus genome^43–46^. A recent first-in-human study in glioblastoma evaluated an Edmonston recombinant measles virus expressing carcinoembryonic antigen ^47^. The trial confirmed the excellent safety of oncolytic MV, its ability to replicate in the tumor, and reported promising, though modest, clinical benefits. Given its capacity to generate DVGs, robustly activate the RIG-I/MAVS pathway, and enhance innate immune and cytolytic responses, MVdeltaC is likely to achieve improved therapeutic efficacy. The enhanced tumor cell killing activity and immunogenicity of MVdeltaC relies on a functional RLR pathway. RIG-I and MDA5 are critical in the body’s antiviral defense, but their roles extend into cancer biology as potential tumor suppressors and immunotherapy targets ^8,48–52^. Their pathways may be exploited to enhance anti-cancer immune responses. RIG-I and MDA5 can both function as tumor suppressors by detecting viral infections and abnormal RNA structures produced by cancer cells, particularly when RNA splicing and editing are disrupted or when endogenous retroviruses are expressed in the form of dsRNA^53^. They trigger an immune response that may inhibit tumor growth or even promote the destruction of cancer cells. MDA5 was originally identified in melanoma, and its activation has shown anti-tumor activity in this type of cancer. Activation of the RLR pathway leads to the production of IFN-I and pro-inflammatory cytokines, thereby promoting an anti-tumor immune response by recruiting immune cells like NK cells and T cells to attack cancer cells^49–52^. Mutations in cancer cells can inhibit RIG-I or MDA5 signaling, allowing tumors to evade immune detection. For instance, some cancers may suppress the expression of IFN-I or downstream signaling proteins to escape immune surveillance, thereby aiding cancer progression^54,55^. RIG-I agonists are currently being explored as therapeutic agents in cancer^56^. These agonists can mimic viral RNA to stimulate an immune response that targets tumors, making RIG-I a focus for cancer immunotherapy. Similarly, activating MDA5 has been shown to enhance checkpoint inhibitor therapies (like anti-PD-1/PD-L1 antibodies), potentially improving the efficacy of immunotherapies in treating various cancers such as melanoma^49^.

In conclusion, we reframed oncolytic MV into an immunotherapeutic virus by genetically disarming it to favor immune stimulation over viral dissemination, a major safety advantage. The unique immunogenic features of MVdeltaC make it a potential candidate for combination with other immunotherapies. Our findings suggest that the anticancer efficacy of MVdeltaC is driven by both direct oncolysis and its capacity to convert the tumor into an immunogenic niche, offering a promising platform for combination with immune checkpoint blockade or cancer vaccines. Indeed, i.t. injections of oncolytic viruses, TLR agonists, or STING agonists are being developed to trigger immune responses by turning “cold” tumors into “hot,” immune-infiltrated ones^57^. MVdeltaC, which is based on a human vaccine with a long-standing an unmatched safety profile, demonstrated by its use for decades for vaccination against measles, is a promising candidate for combined anti-tumor immunotherapeutic strategies. It has now entered GMP manufacturing and is expected to begin Phase I clinical trials soon.

## MATERIALS & METHODS

### Generation of MVdeltaC virus

For MVdeltaC virus cloning, we used the plasmid vector pTM-MVSchw, which contains an infectious MV cDNA corresponding to the anti-genome of the Schwarz MV vaccine strain (GenBank accession no. AF266291.1) and has been described elsewhere^33^. MVdeltaC construct was generated by site-directed mutagenesis on a cassette vector covering the P gene sequence. The second translation initiation codon present in the N-terminal P ORF in the "1 frame was mutated (U -> C), and then a substitution (G -> A) was generated at nucleotide position 1845 of the MV genome to introduce a stop codon within the C ORF. Both mutations do not disturb the P and V ORFs (Fig. 1d). Rescue of MVdeltaC virus was performed using a helper-cell-based system^33^. Briefly, helper HEK293-T7-NP cells were transfected with 5 µg of pTM-MVdeltaC and 0.02 µg of pEMC-La expressing the MV polymerase L gene. After overnight incubation at 37 °C, the transfection medium was replaced by fresh medium, and a heat shock was applied for 3 h at 42 °C and then returned to 37 °C. After two days of incubation at 37 °C, transfected cells were transferred to 100-mm dishes with monolayers of Vero cells (ATCC, CCL-81). Syncytia that appeared after 2–3 days of co-culture were singly picked and transferred onto Vero cells seeded in six-well plates. Infected cells were trypsinized and expanded in 75-cm^2^ and then 150-cm^2^ flasks, in DMEM with 5% Fetal Bovine Serum (FBS, Gibco). To collect viruses, cells were scraped into a small volume of OptiMEM (Thermo Fisher), lysed by a single freeze-thaw cycle and cell lysates clarified by low-speed centrifugation. The infectious supernatant was then collected and stored at −80 °C. Titer of MVdeltaC was determined on Vero cells seeded in 96-well plates infected with serial ten-fold dilutions of virus in DMEM with 5% FBS. After incubation for 7 days, cells were stained with crystal violet, and TCID50 values were calculated using the Karber method^58^. MV-eGFP, MVdeltaC-eGFP, MV-mCherry and MVdeltaC-mCherry were cloned, produced, purified and titrated as previously described^33^.

### Virus growth curves

Monolayers of Vero, HEK293, A549 and HeLa cells in 24-mm-diameter dishes (6-well plates) were infected with MVdeltaC or MV-Schwarz at an MOI of 1. At various times post infection, cells were scraped into culture medium. After freeze-thawing of cells in their medium and clarification of cell debris, virus titers were determined as described above.

### Virus Sequencing

RNA was extracted from the clarified cell supernatants using the QIAamp Viral RNA Kit (Qiagen) following the manufacturer’s instructions. MV genome was reverse transcribed into cDNA and amplified by PCR using previously described primers^33^. Libraries were prepared using a Nextera XT kit (Illumina), before sequencing on an Illumina NextSeq500 (2 × 75 cycles). Raw sequence data were deposited in the Sequence Read Archive (https://www.ncbi.nlm.nih.gov/sra) under BioProject ID PRJNA1309675. For bioinformatic analysis, adaptors and low-quality sequences of raw reads were removed using Trimmomatic v.0.39^59^. We assembled the trimmed reads using megahit v.1.2.9^60^ with default parameters. Raw reads were then mapped on the viral scaffold. The mapping data were visually checked to confirm the accuracy of the genomes using Geneious Prime 2025. Consensus, coverage statistics and intrasample single-nucleotide variants (iSNVs) estimates were obtained using iVar^61^.

### Western blot analysis

Vero cells in six-well plates were infected with MVdeltaC or MV at an MOI of 0.1. At 36–48 h post-infection, infected cells were lysed in RIPA lysis buffer (Thermo Fisher). Samples were briefly centrifuged and subjected to 4–12% gradient on a NUPAGE-PAGE gel (Invitrogen). After transfer to a nitrocellulose membrane (GE Healthcare), the membrane was subsequently probed with a rabbit polyclonal anti-MV C antibody (Pr Takeuchi, Tsukuba University, Japan, 1:2000 dilution) followed by a horseradish peroxidase (HRP)-conjugated swine anti-rabbit IgG antibody (P0399, Dako, 1:3000 dilution). Bands were visualized using SuperSignal West Pico Plus chemiluminescent HRP substrate (Thermo Fisher). For loading controls, membranes were stripped with 5% NaOH for 5 min and re-probed with a mouse monoclonal anti-MV-N antibody (ab9397, Abcam, 1:20,000 dilution) followed by an HRP-conjugated anti-mouse IgG (NA931V, GE Healthcare, 1:10,000 dilution).

### Cell culture

Human mesothelioma cell lines (Meso 4, Meso 11, Meso 13, Meso 34, Meso 35, Meso 47, Meso 56, Meso 76, Meso 150, Meso 163, Meso 173 and Meso 225) were established from pleural effusions and genetically characterized (Biocollection DC-2011–1399, CHU Nantes, France) ^62^. The human melanoma cell line M6 was a kind gift from Nathalie Labarrière (INCIT, biocollection PC-U892-NL, CHU Nantes, France). The cell lines were cultured at 37°C and 5% CO_2_ in Roswell Park Memorial Institute Medium (RPMI)-1640 supplemented with 2 mM L-Glutamine, 100 IU/mL Penicillin, 100 µg/mL Streptomycin (all from Gibco), and 10% heat-inactivated FBS (Corning). Normal peritoneal mesothelial cells MES-F and LP-9 were purchased from Tebu-bio and the Coriell Institute, respectively, and cultured in their specific medium according to the manufacturers’ recommendations. Mouse neuroblastoma cell line NS20Y (DSMZ# ACC 94), A549 cells human adenocarcinoma alveolar basal epithelial cells, and HeLa cervical cancer cells were cultured in Dulbecco’s modified Eagle’s medium (DMEM) supplemented with 10% of heat-inactivated FBS, and 100 U/ml penicillin, 100 U/ml streptomycin (Thermo Fisher) in a 5% CO_2_ atmosphere. Cells were routinely checked for Mycoplasma contamination (PlasmoTest, InvivoGen).

### Virus infection of mesothelioma cells

MV and MVdeltaC infections were analyzed by time-lapse fluorescence microscopy. One day before infection, healthy cells and mesothelioma cell lines were seeded in 24-well plates at a density of 1.5 × 10^5^ cells per well. Cells were infected with MV-eGFP or MVdeltaC-eGFP (MOI 1) and images were acquired every 10 minutes for 3 to 4 days with a Leica DMI6000B microscope using a 10X objective and the MetaMorph® Microscopy Automation & Image Analysis Software (version 7.8) at the MicroPICell core facility (Nantes, France). Subsequent analyses were performed with the Fiji software^63^.

### Cell viability and cell death analyses

Cell viability was assessed using the CellTiter-Glo Luminescent Cell Viability Assay (Promega, Cat# G7571), according to the manufacturer’s instructions. Briefly, cells were seeded at a density of 4 × 10^4^ cells per well in 50 µL of culture medium in white, flat-bottom 96-well plates (Corning). Virus stocks were diluted to achieve MOIs of 1, 5, or 10, and 50 µL of the viral dilution was added to each well immediately after seeding. Infections were performed in triplicate. Plates were incubated at 37 °C with 5% CO_2_ for 3–6 h to allow viral entry, followed by medium replacement to remove residual inoculum. At 24 to 96 h post-infection, 100 µL of culture supernatant was removed, and 100 µL of CellTiter-Glo reagent was added directly to the wells. Luminescence was recorded using an EnSpire Multimode Plate Reader (PerkinElmer).

Caspases-3/7 activation was measured in infected cells by either luminescence assay or fluorescence microscopy. For the luminescence assay, mesothelioma cells were seeded in a 96-well plate at a density of 15,000 cells per well and cultured for 24h. Cells were then infected with either MV or MVdeltaC as described above. Caspases-3/7 activity was quantified daily from day 2 to day 6 using the Caspase-Glo® 3/7 assay kit (Promega). Briefly, 100 μl of Caspase-Glo® reagent was added onto the cells and incubated at room temperature for 1h. The luminescent signal of each sample was then measured for 1 second using a Mithras LB 940 luminometer (Berthold Technologies).

For the microscopy experiments, cells were seeded in a 96-well plate at a density of 50,000 cells per well. After 24h, cells were infected with either MV-mCherry or MVdeltaC-cherry as described above. Viral inoculum was replaced by fresh culture medium mixed with NucView® 405 caspase-3 substrate (Ozyme) at a final concentration of 1µM. Images were acquired every day for 4 days with an Axio Observer Z1 microscope and the ZEN software (Carl Zeiss). Images were further analyzed with the Fiji Software.

Quantities of HMGB1 protein released from dying cells were measured by ELISA (IBL International) according to the manufacturer’s instructions 72h after infection of 0.5 × 10^6^ mesothelial or mesothelioma cells with MV or MVdeltaC (MOI 1) in 6-well plates.

### dsRNA immunofluorescence staining

One day before infection, PM cells were seeded in µ-Slides 8-well plates (ibidi) at a density of 40,000 cells per well. Cells were infected with either MV-eGFP or MVdelta-eGFP (MOI 1). dsRNA staining was performed at 48h post-infection. Briefly, cells were fixed with PBS containing 4% paraformaldehyde (PFA, Electron Microscopy Sciences) for 5 min, followed by PBS washes. Cells were then permeabilized with PBS containing 0.1% Triton X-100 (Sigma-Aldrich) for 10 min. Plates were incubated with an anti-dsRNA antibody (clone J2, English & Scientific Consulting) diluted in PBS containing 0.1% Triton X-100 for 1h at room temperature, followed by 2 PBS washes. Cells were then incubated with an Alexa Fluor 647-coupled anti-mouse secondary antibody (Molecular Probes, Thermo Fisher Scientific) for 20 min at room temperature. The slides were then washed twice with PBS. Nuclei were stained with Hoechst (Thermo Fisher Scientific) for 5 min. Finally, cells were fixed for 20 min in PBS 4% PFA and then stored in PBS 0.5% PFA until analysis with a Nikon A1RS fluorescence confocal microscope (MicroPICell core facility).

### RNA extraction and quantitative real-time PCR in HAP cells

HAP, HAP RIG-I KO, HAP MDA5 KO, and HAP MAVS KO cells were infected at an MOI of 1 with MV Schwarz, MVdeltaC, or left uninfected (mock), and incubated at 37°C in a humidified atmosphere containing 5% CO_2_. Total RNA was extracted 24 hours post-infection using the RNeasy Mini Kit (QIAGEN) according to the manufacturer’s instructions. RNA concentration and purity were assessed using a NanoDrop spectrophotometer (Thermo Fisher Scientific). Expression levels of selected genes were quantified using one-step real-time PCR with the TaqMan RNA-to-Ct 1-Step Kit (Thermo Fisher), following the manufacturer’s protocol. One hundred nanograms of total RNA were used per reaction with 20X TaqMan Gene Expression Assays (*CXCL10*: Hs00171042_m1; *TNF*: Hs00174128_m1; *MX1*: Hs00895608_m1; *CD274*: Hs00204257_m1; *TNFRSF10B*: Hs00366278_m1; *MICA*: Hs00792195_m1; *ULBP2*: Hs00607609_m1; *GAPDH*: Hs02786624_g1; *IFNB1*: Hs01077958_s1) (Life Technologies). All reactions were performed in triplicate on the QuantStudio™ 6 Flex Real-Time PCR System (Applied Biosystems). Target gene expression levels were normalized to the endogenous control GAPDH, and relative expression was calculated using the comparative Ct method (ΔΔCt).

### Virus Sequencing from infected HAP cells

HAP cells were infected at an MOI of 1 with MV Schwarz, MVdeltaC, MV 2N (MV Schwarz virus expressing two copies of the N protein; used as a positive control) or left uninfected (mock). Cells were incubated at 37 °C in a humidified atmosphere containing 5% CO_2_. Total RNA was extracted 24 hours post-infection using the RNeasy Mini Kit (QIAGEN) according to the manufacturer’s instructions. RNA quality and integrity were assessed using the Agilent RNA 6000 Nano Kit (Agilent Technologies) on the Agilent 2100 Bioanalyzer system. The RNA Integrity Number (RIN) was used as a quality control metric for downstream applications. Libraries were prepared from 150 ng of RNA using the Illumina Stranded Total RNA Prep with Ribo-Zero Plus kit (Illumina), according to the manufacturer’s protocol, with 13 PCR cycles. Unbound adaptors and primers were removed by two successive purifications using AMPure beads (Beckman Coulter) at a 0.8:1 bead-to-sample ratio. The resulting libraries contained fragments ranging from 200 to 1000 bp, with a peak around 450 bp, as determined on a 5300 Fragment Analyzer (Agilent Technologies). Then, libraries were pooled, diluted to 0.75 nM, and sequenced on a NextSeq 2000 system (Illumina) with a P1 100-cycle kit to generate 58-base paired-end reads (40–50 million reads per sample).

### Defective interfering RNA genomes (DI RNA) bioinformatic analysis

To identify potential DI RNA, merged FASTQ files were analyzed using the Python script, DI-tector (http://www.di-tector.cyame.eu/)^36^, with the minimum segment length set to 25. Reads that aligned perfectly to the viral reference genome (GenBank accession number no. FJ211590) were excluded. Reads aligning to two separate parts of the viral genome on either side of a junction, indicative of a breakpoint and reinitiation site, were then extracted and classified as DVGs.

### Co-culture of infected tumor cells with human monocyte-derived dendritic cells

Human PBMCs were obtained from platelet apheresis residues of healthy donors after informed consent (Etablissement Français du Sang, Nantes, France) under the agreement NTS-2014–06. After Ficoll density gradient centrifugation (Eurobio-AbCys), monocytes were purified from PBMCs by counterflow centrifugation elutriation using a Beckman Avanti J20 centrifuge equipped with a JE5.0 rotor and a 7 mL elutriation chamber (DTC core facility, CIC-BT-0503, Nantes, France). Monocyte purity (> 85%) was assessed by flow cytometry. Monocytes were differentiated into DCs in 6-well plates at 2 × 10^6^ cells/mL in RPMI-1640 supplemented with 2 mM L-glutamine, 100 U/mL penicillin, 100 μg/mL streptomycin and 2% (v/v) human albumin (Laboratoire Français de Fractionnement et de Biotechnologies), 1,000 U/mL recombinant human GM-CSF and 200 U/mL recombinant human IL-4 (both from CellGenix). Immature DCs (iDC) were harvested at day 6 and cultured for 24h at 1 × 10^6^ cells/mL in 24-well under different treatment conditions. After 24h, DC maturation was assessed by flow cytometry (FACS Canto II, CytoCell core facility, Nantes, France) after double staining with a PE-conjugated anti-HLA-DR antibody (clone AC122, Miltenyi Biotec) and a BV421-conjugated anti-CD83 antibody (clone HB15e, BD Biosciences). The percentage of matured DCs was determined as CD83^+^ cells among all HLA-DR^+^ cells. All flow cytometry analyses were performed using the FlowJo software (version 7.6.5, FlowJo LLC).

### NOD SCID mice studies

Six-week-old female NOD SCID mice were obtained from Charles River Laboratories. Mesothelioma xenograft models were established by challenging the mice intraperitoneally with 5 × 10^6^ Meso 34 or Meso 163 cells. Three weeks or two months post-challenge for Meso 163 and Meso 34, respectively, mice were randomized into three groups and injected intraperitoneally with 100 µL of control saline (PBS), MV-eGFP (2 × 10^5^ TCID_50_) or MVdeltaC-eGFP (2 × 10^5^ TCID_50_). Fifteen days after treatment, animals were sacrificed, and metastatic nodules or residual tumor tissues were collected and fixed in 4% paraformaldehyde (Electron Microscopy Sciences) for weighing. The M6 melanoma xenograft model was established by challenging the mice subcutaneously with 1 × 10^6^ cells. Mice were injected intratumorally twice at days 10 and 17 post-challenge with control saline (PBS), MV or MVdeltaC. Tumor growth was measured every 4-5 days with a digital caliper and tumor volumes were calculated according to V = (length × (width)^2^/2. For histological examination, the paraformaldehyde fixed, paraffin-embedded sections of tumors and surrounding invaded tissues, when present, were cut with a Bond Max automaton (Menarini) and stained with hematoxylin-phloxine-saffron (HPS). Slides were scanned with a Nanozoomer 2.0 HT (Hamamatsu Photonics). These protocols were approved by the local ethical committee (CEEA PdL n°6) and the French Ministry of Research (#01257.03).

### PDX studies

The primary human mesothelioma PDX models were derived from patients with pleural or peritoneal mesothelioma and used as xenotransplantation model in immunodeficient female NMRI:nu/nu mice. The models were established by the EPO Berlin-Buch GmbH. The tumors were propagated *in vivo* and tumor tissue from one *in vivo* passage were used for s.c. implantation in the inguinal region of female nude mice. After an acclimatization time of 4 days, tumor fragments (2x2mm / mouse) from an *in vivo* passage were transplanted s.c. into 13 female nude mice (day 0). For tumor transplantation the animals were anesthetized by a single intravenous injection (0.10 mL / mouse) with Propofol-® Lipuro (0.5 mg / mouse). The skin of the mice was disinfected with isopropanol (70 % v/v). A superficial vertical incision in the skin of 5-6 mm on the left flank was performed. The tip of a surgical scissor was inserted into the incision and was used to form a pocket in the subcutaneous space. One tumor fragment per mouse was implanted into the pocket using surgical tweezers. Finally, the incision was closed with a metal clip. Studies were performed under Biosafety level S2. After the xenotransplantation, the engraftment and the propagation of the tumor in the mice were controlled at least twice weekly by palpation. When the tumor was palpable, the measurements of tumor diameters were performed with a digital caliper twice weekly. Prior starting the treatment, animals were randomly assigned into treatment and control groups (n=3). During randomization, the animals were individually labeled by earmarks. Animals were treated i.t. with MVdeltaC at doses 1 × 10^5^ and 1 × 10^6^ TCID_50_ per animal once weekly for up to 10 weeks. Virus was injected i.t. with three injections of 30-35µl at different sites in the tumor. Animals from the control group received the vehicle also i.t. Tumor diameters were measured three times weekly with a caliper. Tumor volumes were calculated according to V = (length × (width)^2^)/2. For calculation of the relative tumor volume (RTV) the volumes at each measurement day are related to the day of first treatment. These RTV values were used for the evaluation according to the modified RECIST criteria, with RTV values > 1.3 as Progression (P), RTV between 0.7 and 1.3 as Stable Disease (SD), RTV < 0.7 as Partial Regression (PR) and RTV = 0 as complete Regression (CR). At each measurement day, the median and mean tumor volumes per group and the treated to control (T/C) values in percent were calculated. The experiment was terminated when tumor volumes reached a weight of >10% of the animal’s normal body weight (typically a mean s.c. flank diameter of 17 mm or a tumor volume >1.5cm³ in a 25 g mouse). Body weight was determined 3 times weekly and all animals were checked daily for treatment-related toxicity. The decrease in body weight provided a measure of treatment-related toxicity.

### Syngeneic A/J mouse model

Six- to 8-week-old female A/J mice were purchased from Envigo (Inotiv). Animals were housed in the Institut Pasteur BIME A3 animal facility on a 12 h light/dark cycle at 18–24°C and humidity of 40-60%. All the experiments were performed in class III safety cabinets in the Institut Pasteur animal facilities accredited by the French Ministry of Agriculture for performing experiments on live rodents. Animal experiments were validated by Institut Pasteur ethics committee with the approval numbers: APAFIS #35041-2022013107218823 v5, dap210133 and CHSCT 16-013. All animals were handled in strict accordance with good animal practice.

#### Tumor transplantations

A total of 5 × 10^6^ NS20Y cells in 100µL of PBS were injected subcutaneously in the right shaved flank of the mice. Tumor sizes were monitored with a digital caliper (Fisherbrand) every 2 to 3 days and tumor volume was calculated using formula V = L x W^2^ x 0.5, where V is the tumor volume, L is the tumor length, and W is the tumor width. Health monitoring and humane endpoints were implemented using Institut Pasteur ethics committee validated health evaluation scoring grid. Animals were killed when tumor volume exceeded ≥1500mm^3^, if the tumors developed hemorrhagic ulcers, if a body weight loss ≥20% occurred, if mice showed bad general condition scores, or the purpose of the experiment has been reached.

#### Tumor immunotherapy treatment

Treatments started on day 6 after tumor cell injection when tumors reached 100-250mm^3^ (7 to 10mm in largest diameter). Intratumoral injections of MVdeltaC in 100µl volume were performed on days 6, 9 and 12 after NS20Y cells injection. The 100µL dose of PBS or MVdeltaC was injected in one shot during the first treatment and using the “radial technique” during next treatments. In “radial technique” a single-entry point in the punctured tumor was performed to minimize discharge and bleeding. The needle was then gently moved within the tumor mass to reach and inject different tumor parts. The immune checkpoint-targeted antibody anti-CTLA-4 (clone 9H10, BioXCell) was injected systemically via the intraperitoneal route at the concentration of 200µg in 100µl of PBS, four times on days 6, 9, 12, and 15 after NS20Y cells injection.

#### CD4/CD8 T and NK cell depletions

Anti-CD4 (rat IgG2b; clone GK1.5), anti-CD8a (rat IgG2b; clone 2.43), and anti-CD20 (rat IgG2c; clone MB20-11) depleting mAbs were purchased from BioXCell. NK cells depleting anti-asialo GM1 antibody (polyclonal rabbit anti-mouse IgG) was purchased from Thermo Fisher. Depleting antibodies were injected intraperitoneally 1 day before starting the intratumoral MVdeltaC therapy. To deplete T cells, anti-CD4 or anti-CD8 mAbs (200 µg in 100 µL of PBS) were injected intraperitoneally on days 5, 6 and 7 after NS20Y cells injection. Targeted depletion of NK or B cell was performed on days 5 and 7 following NS20Y tumor cell inoculation using 50µg of anti-asialo GM1 or 200µg of anti-CD20 antibodies, respectively. Sustained immune cell depletion was achieved by weekly antibody administrations starting from day 7 until day 49. Depletion efficacy was evaluated in blood samples of selected mice by flow cytometry. *Flow cytometry.* Immune cell depletion efficacy was evaluated in the blood from three mice of each group. Blood was collected in Microvette 500 EDTA K3E tubes (Sarstedt) by submandibular vein puncture. Cells in the blood samples (200 µL) were sedimented by centrifugation, supernatant was discarded and red blood cells were lysed using RBC lysis buffer (Sigma-Aldrich). Cells were washed with PBS 2% FBS buffer and surface-stained with eBioscience CD3e (clone 145-2C11), CD4 (clone RM4- 5), CD8a (clone 53-6.7), CD335 (NKp46, clone 29A1.4), and Biolegend CD19 (clone 6D5) specific monoclonal antibodies. The stained cells were acquired on an Attune™ NxT Flow Cytometer (Thermo Fischer).

#### Ethics

All animal experiments were performed according to French legislation in compliance with the European Communities Council Directives (2010/63/UE, French Law 2013–118, 6 February 2013) and according to the regulations of Institut Pasteur Animal Care Committees. The Animal Experimentation Ethics Committee (CETEA 89) of the Institut Pasteur approved this study (dap210133) before experiments were initiated.

#### Immunohistochemistry

On day 13, tumors were harvested and fixed in 10% formol for 48h. Tumors were photographed and embedded in the Optimal Cutting Temperature compound (Tissue-Tek Sakura). Immunohistochemical analysis of spleen and tumor 8μm-thick cryostat sections was performed on Leica Bond RX using rabbit anti-mouse CD4 (Cell signaling, D7D2Z) or CD8a (Cell signaling, D4W2Z) monoclonal antibodies, followed by biotinylated goat anti-rabbit secondary antibody (Dako) and streptavidin-HRP, provided in the Leica Bond RX kit. Slides were scanned using an Axioscan Z1 Zeiss slide scanner and the images were analyzed using Zen 3.8 software.

## Statistical analysis

Analyses were performed with the GraphPad Prism software (versions 6 to 10, GraphPad Software Inc., La Jolla, CA, USA). Unpaired, non-parametric, Mann-Whitney was used to compare two independent groups. Unpaired, non-parametric, Kruskal-Wallis was used to compare three or more independent groups. Differences were considered significant when *p < 0.05, **p < 0.01 or ***p < 0.001. All data are from at least three independent experiments and are presented as mean ± standard deviation (SD).

## DATA AVAILABILITY

The authors declare that the data supporting the findings of this study are available within this article and its supplementary information files or available from the corresponding author upon reasonable request. The source data files are provided with this paper.

## ACKNOWLEDGMENTS

This work was funded by Oncovita and by grants from the Agence Nationale de la Recherche (ANR-16-CE18-0016), the Fondation pour la Recherche Médicale (Labellisation DEQ20140329511) and La Ligue Contre le Cancer Grand Ouest. We thank the staff of the animal facility of the Pasteur Institute for mouse care, and the Cytocell - Flow Cytometry and FACS core facility (SFR Bonamy, BioCore, Inserm UMS 016, CNRS UAR 3556, Nantes, France) for its technical expertise and help, member of the Scientific Interest Group (GIS) Biogenouest and the Labex IGO program supported by the French National Research Agency (n°ANR-11-LABX-0016-01). We acknowledge Philippe Hulin, Steven Nedellec and Stéphanie Blandin from the MicroPICell core facility (SFR Bonamy, BioCore, Inserm UMS 016, CNRS UAR 3556, Nantes, France), member of the Scientific Interest Group (GIS) Biogenouest, IBISA, and the national infrastructure France-Bioimaging supported by the French National Research Agency (ANR-24-INBS-0005 FBI BIOGEN). We acknowledge Elodie Turc and Laure Lemée, Biomics Platform, C2RT, Institut Pasteur, Paris, France, supported by France Génomique (ANR-10-INBS-09) and IBISA. We extend special thanks to Delphine Coulais and Sylvia Lambot for their assistance regarding monocyte purification and animal experiments, respectively.

## AUTHOR CONTRIBUTIONS

AB designed and performed most experiments and contributed to writing the manuscript. HVP, JSN, VN, CC, AA, SG, PNF, RV, ES, MC, DP, TP performed in vitro and in vivo experiments. MP and ESL performed NGS analyses. DH and SL performed histological analyses. BB and JH performed PDX experiments. VR and JFLB contributed to preclinical design. MG, JFF and AVK advised and assisted in proofreading the manuscript. AB, NB and FT wrote the manuscript. FT conceived and supervised the overall project and heads the laboratory.

---

**Figure S1.**
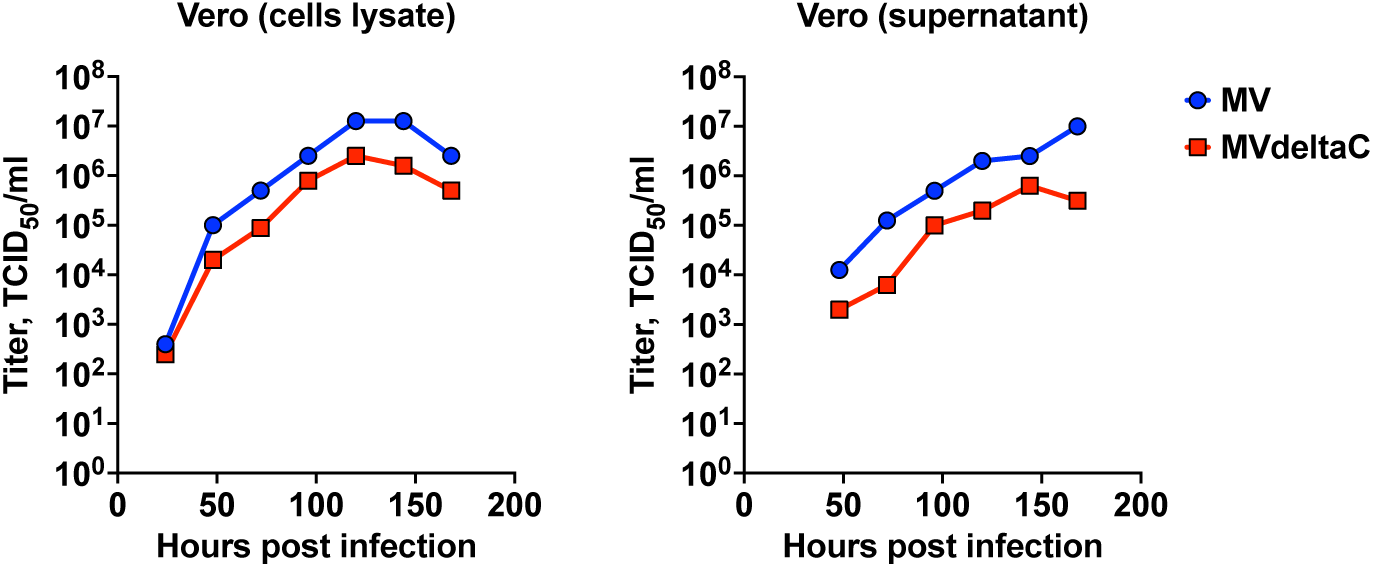
Growth kinetics of MV and MVdeltaC on Vero cells. Cells were infected at MOI 0.01 and incubated at 32°C. The data show reciprocal endpoint dilution titers of cell-associated (left) and cell-free (right) viruses.

**Figure S2.**
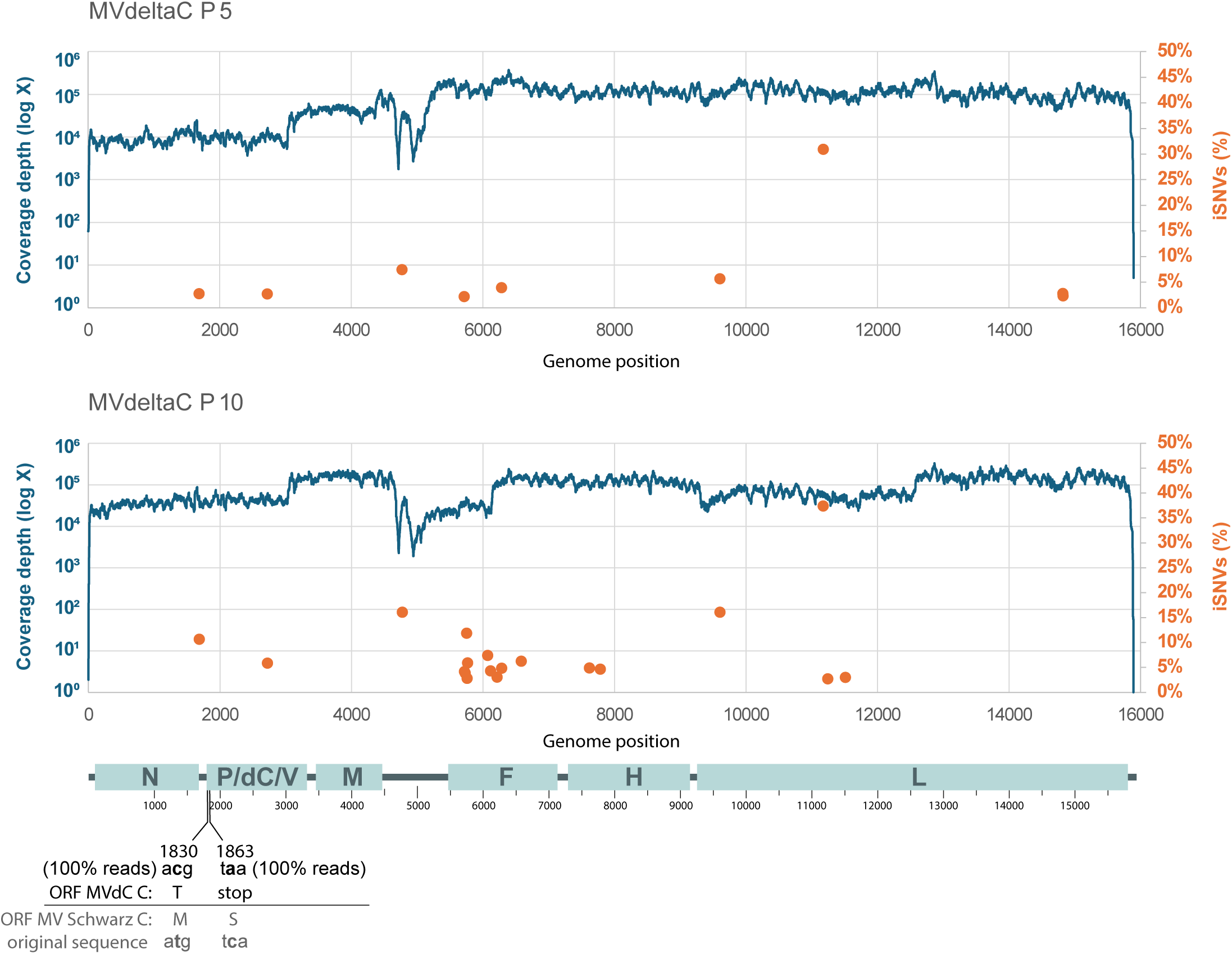
Sequencing analysis of MVdeltaC passages. Reads coverage and proportion of intrasample single nucleotide variants (iSNVs) obtained from next-generation sequencing of MVdeltaC at passages 5 (P5) and 10 (P10). A schematic of the measles virus genome is shown below using the same scale, with the two engineered mutations preventing C protein expression highlighted. No sequence changes were detected at either of these positions in P5 or P10 samples.

**Figure S3.**
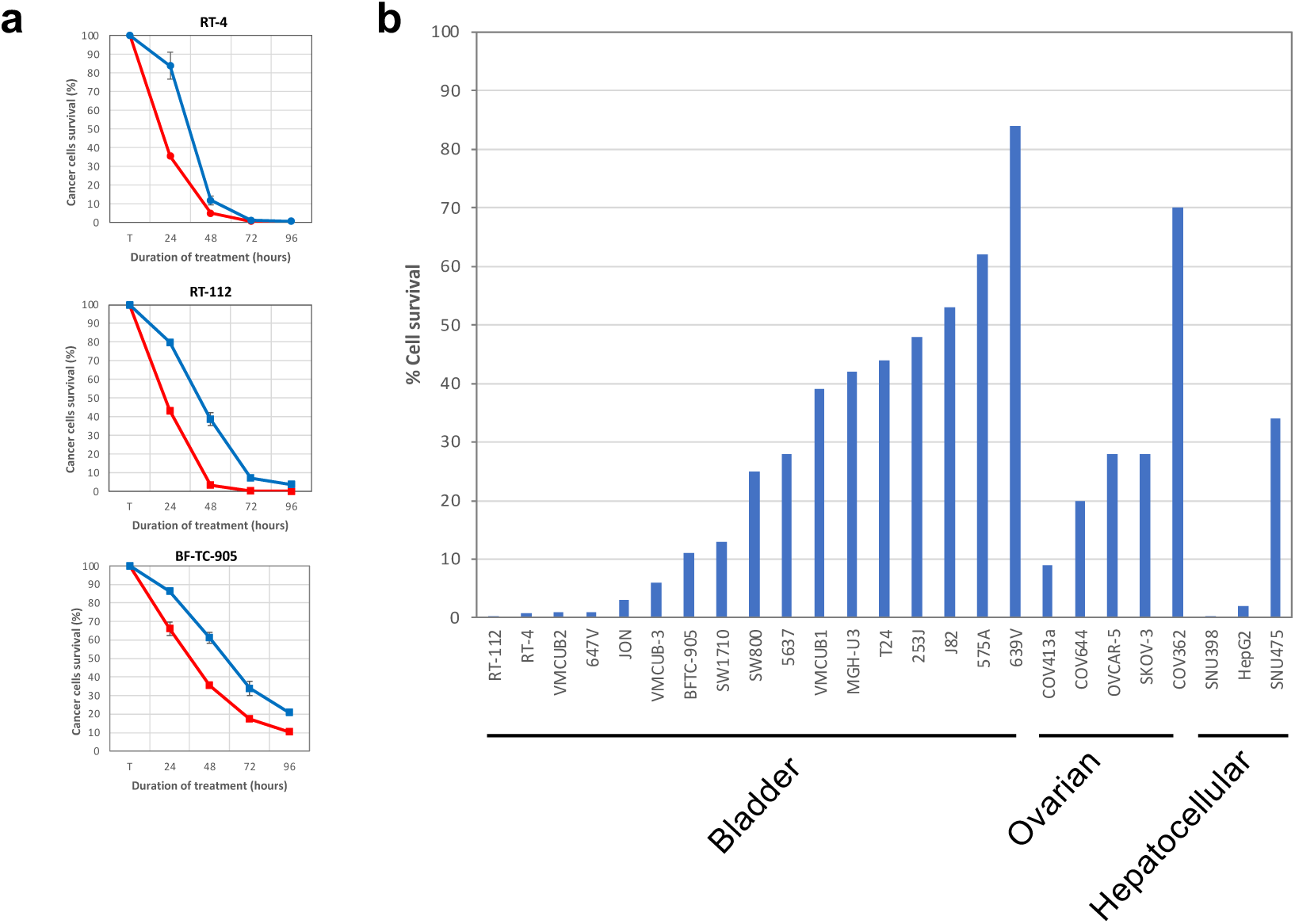
Infection and killing of human cancer cells by MVdeltaC. (a) Cell survival of three human bladder cancer cell lines after infection with MV (blue) or MVdeltaC (red) at MOI 5, measured by CellTiter-Glo assay; means ± SD; measurements are in triplicate; data normalized to non-infected cells. (b) MVdeltaC killing of a series of human cancer cells of bladder, ovarian or hepatocellular origin as determined by CellTiter-Glo assay (cancer cells survival after 72 hours of infection at MOI 5).

**Figure S4.**
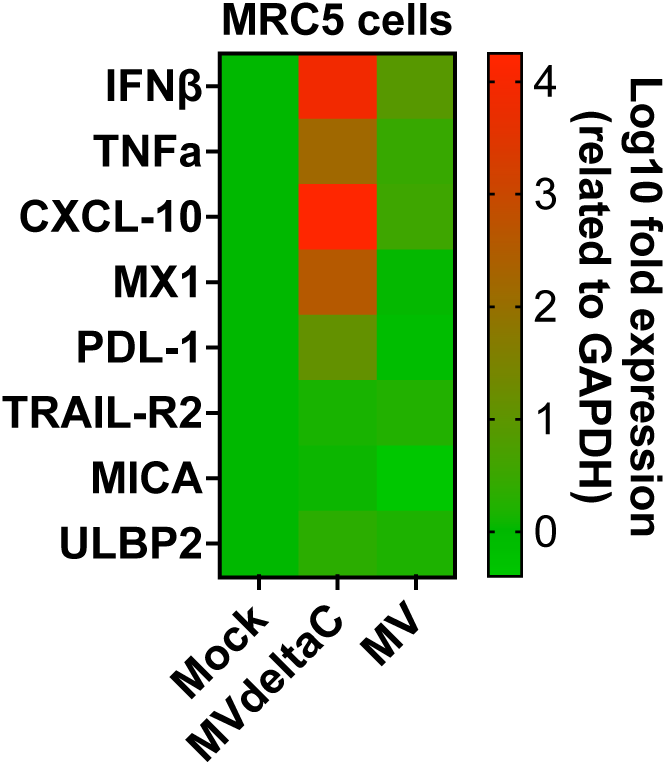
Gene expression analysis in infected MRC5 cells. Heatmap of differentially expressed genes in MRC5 cells after 24h incubation with MV, MVdeltaC (MOI 1) or PBS (Mock); experiment in triplicate, a gradient of colors from green to red represent log_10_ fold change values from lowest to highest, respectively.

**Figure S5.**
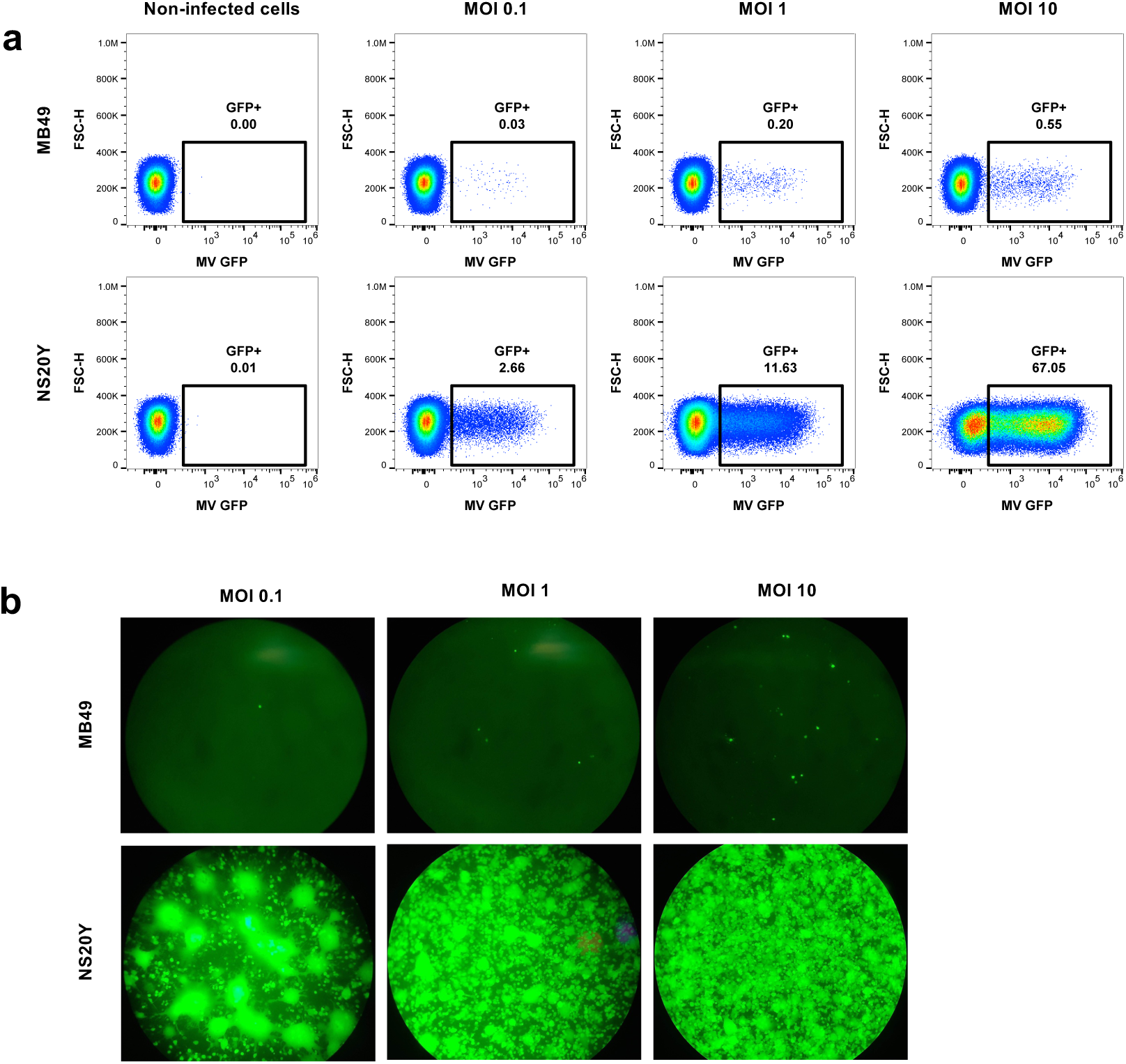
NS20Y cells are permissive to measles infection. Mouse neuroblastoma NS20Y cells and bladder carcinoma MB49 cells were infected with MV-eGFP at the indicated MOI and analyzed by (a) flow cytometry 24 hours post-infection, and (b) immunofluorescence microscopy 72 h hours post-infection.

**Figure S6.**
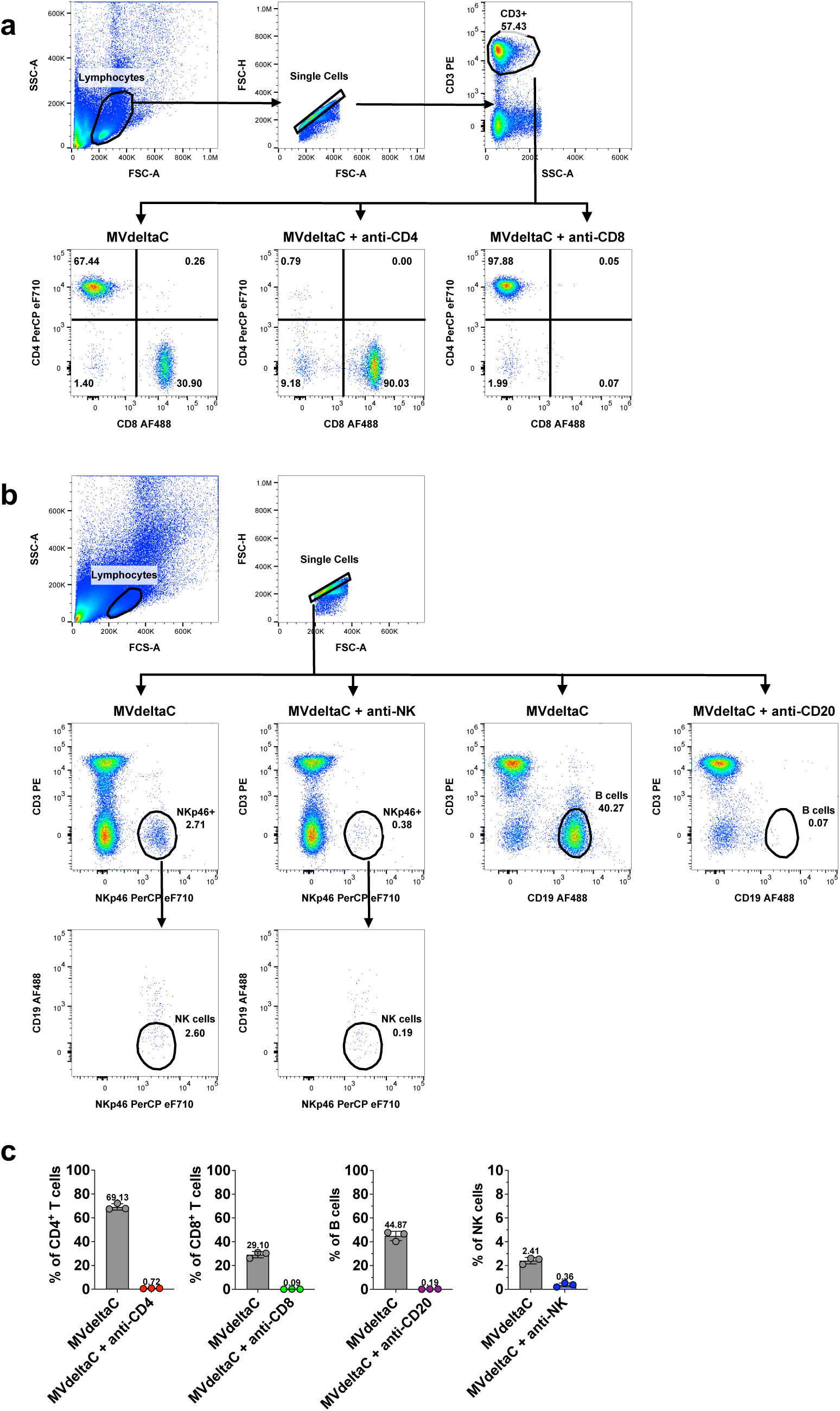
Flow cytometry analysis of immune cell populations in the blood of A/J mice depleted for CD4^+^, CD8^+^, B or NK cells. (a) Representative dot plots show the flow cytometry gating strategy to identify CD4^+^ (CD3^+^ CD4^+^ CD8^-^), CD8^+^ (CD3^+^ CD8^+^ CD4^-^) or (b) NK (CD3^-^NKp46^+^ CD19^-^) and B (CD3^-^ CD19^+^) cell populations in the blood of treated mice. Mice were grafted with 5x10^6^ NS20Y murine tumor cells on day 0. T cell depletion was achieved by i.p. injection of anti-CD4 or anti-CD8 mAbs (200 µg in 100 µL of PBS) on days 5, 6 and 7 after NS20Y cells injection. NK or B cell depletion was performed on days 5 and 7 using 50µg of anti-asialo GM1 or 200µg of anti-CD20 antibodies, respectively. Sustained immune cell depletion was achieved by weekly antibody administrations on days 14 and 21. Tumor therapy with MVdeltaC at 2x10^6^TCID_50_ was performed on days 6, 9 and 12. Immune cell populations in the groups of A/J mice were analyzed on day 27. (c) Bar graphs show flow cytometry data for the three animals from each group, means ± SD.

**Figure S7.**
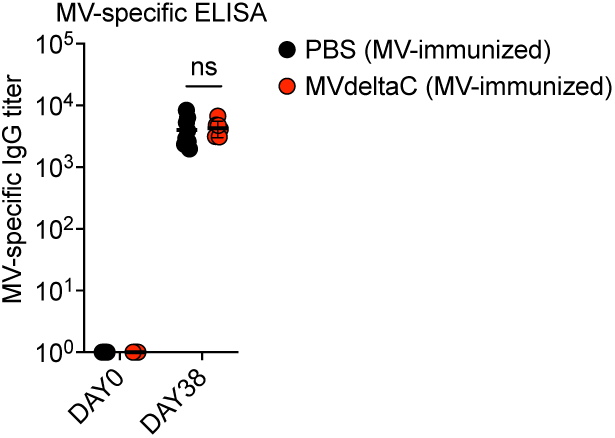
MV-specific IgG antibody responses in A/J mice following MV Schwarz vaccination. Mice were immunized i.p. with MV Schwarz (1 × 10^5^ TCID_50_) on day 0. Sera were collected at day 38 post-immunization and assessed for IgG antibody responses to MV antigens by ELISA. The data show the reciprocal endpoint dilution titers of MV-specific IgG antibodies in preimmune (day 0) and day 38 sera; each point represents an individual mouse; means ± SD, n= 7-9 per group; unpaired two-tailed t test; ns, not significant.

**Figure S8.**
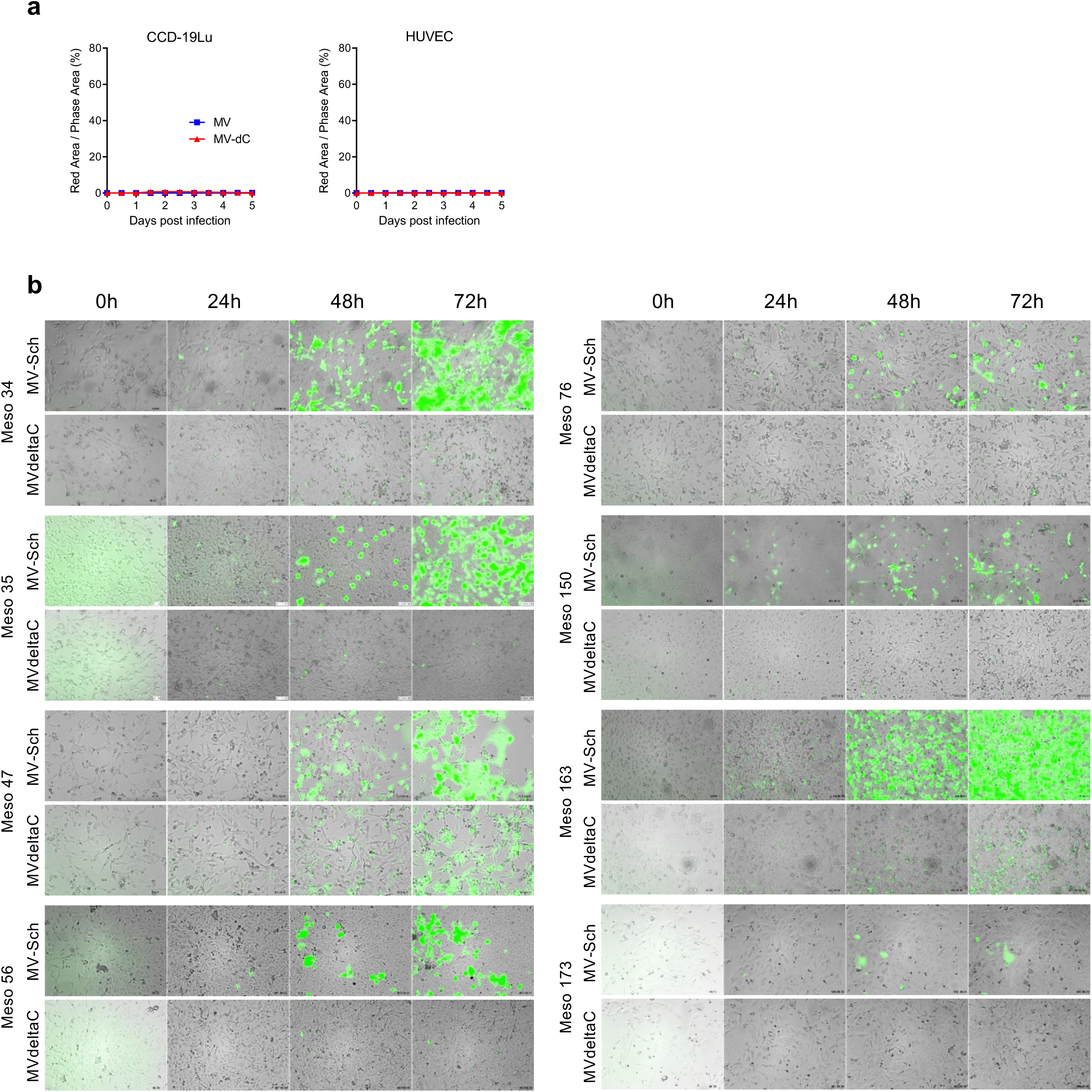
Infection of healthy and cancer human cells by MV or MVdeltaC. (a) Quantification of virus propagation in healthy lung fibroblasts (CCD19-Lu) or endothelial cells (HUVECs) infected with either MV-mCherry or MVdeltaC-mCherry. (b) Time-lapse microscopy analysis of mesothelioma cell lines infected with MV-eGFP or MVdeltaC-eGFP (MOI = 1).

**Figure S9.** Infection of Meso 13 mesothelioma cells by MV or MVdeltaC. Time-lapse microscopy analysis of mesothelioma cell lines infected with MV-eGFP (a) or MVdeltaC-eGFP (b) (MOI = 1).

**Figure S10.**
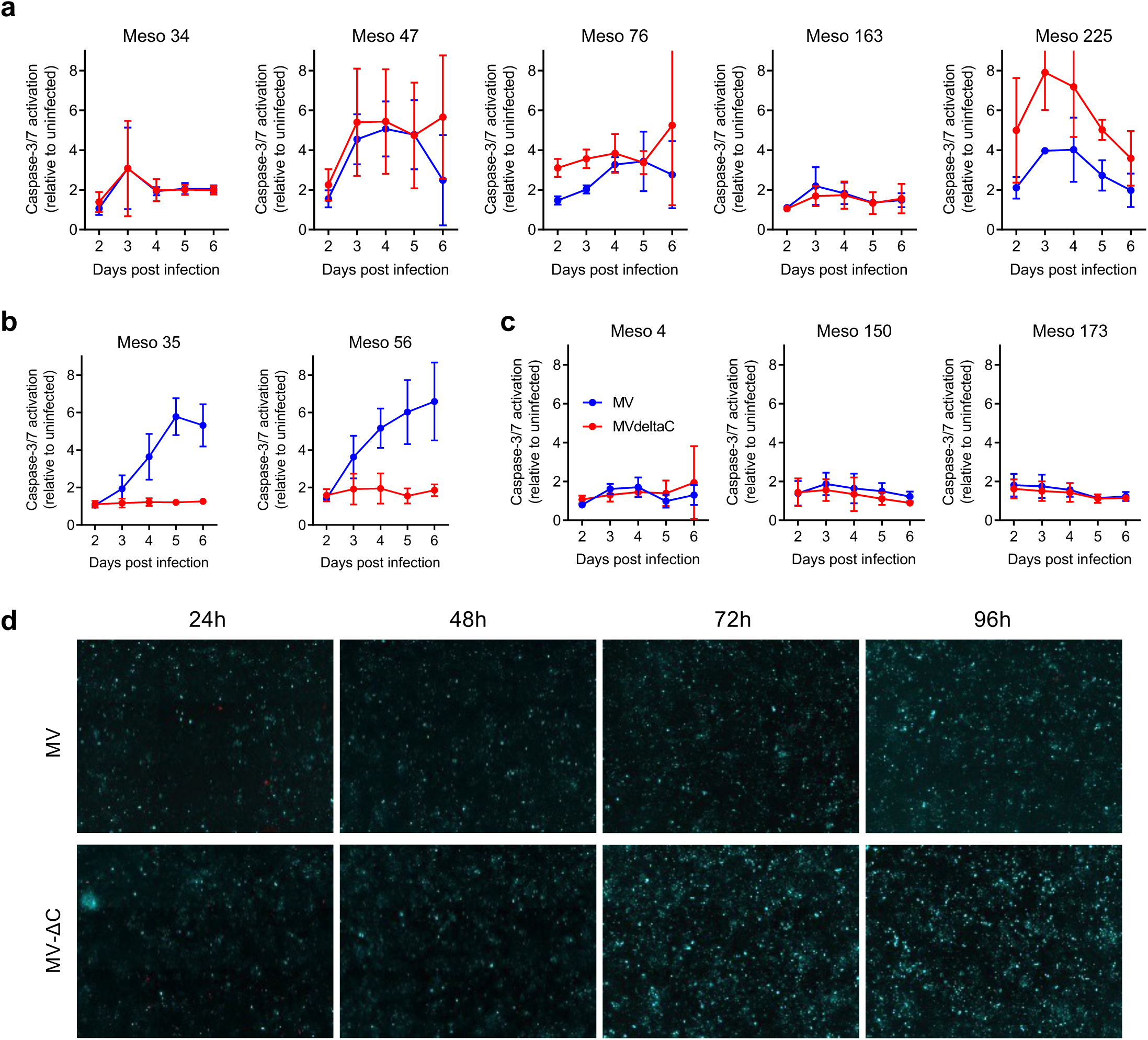
Activation of Caspases-3/7 in mesothelioma cells infected by MVdeltaC. (a-c) Luminescence-based quantification of caspase-3/7 activity in human mesothelioma cell lines. (a) MV-permissive, (b) MVdeltaC-resistant, and (c) MV-resistant cells. (d) Fluorescence microscopy of Meso 4 cells showing caspase-3/7 activation (cyan) after infection with MV-mCherry or MVdeltaC-mCherry (red).

**Figure S11.**
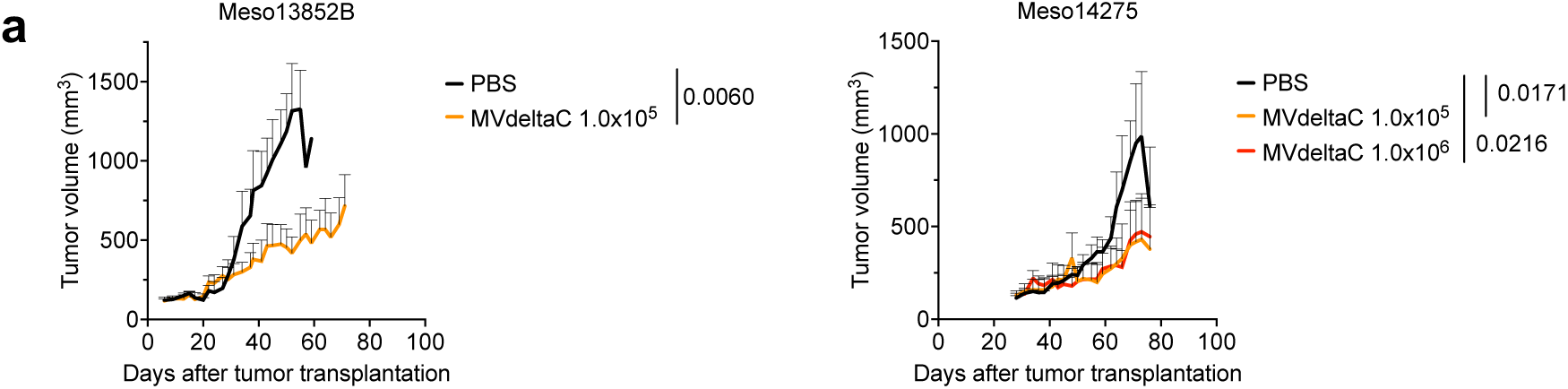
**MVdeltaC inhibits human PDX growth in immunodeficient mice**. Growth of mesothelioma PDX tumors in NMRI nu/nu mice after weekly MVdeltaC therapy at two doses; means ± SEM, n = 3 mice per group; each group was compared to PBS group (unpaired one-tailed t test).

